# Chromosome-level Assemblies of Three *Candidatus* Liberibacter solanacearum Vectors: *Dyspersa apicalis* (Förster, 1848)*, Dyspersa pallida* (Burckhardt, 1986), and *Trioza urticae* (Linnaeus, 1758) (Hemiptera: Psylloidea)

**DOI:** 10.1101/2024.12.03.626329

**Authors:** Thomas Heaven, Thomas C. Mathers, Sam T. Mugford, Anna Jordan, Christa Lethmayer, Anne I. Nissinen, Lars-Arne Høgetveit, Fiona Highet, Victor Soria-Carrasco, Jason Sumner-Kalkun, Jay K. Goldberg, Saskia A. Hogenhout

## Abstract

Psyllids are major vectors of plant diseases, including *Candidatus* Liberibacter solanacearum (CLso), the bacterial agent associated with ‘zebra chip’ disease in potatoes and ‘carrot yellows’ disease in carrot. Despite their agricultural significance, there is limited knowledge on the genome structure and genetic diversity of psyllids. In this study, we provide chromosome-level genome assemblies for three psyllid species known to transmit CLso: *Dyspersa apicalis* (carrot psyllid), *Dyspersa pallida*, and *Trioza urticae* (nettle psyllid). As *D. apicalis* is recognised as the primary vector of CLso by carrot growers in Northern Europe, we also resequenced populations of this species from Finland, Norway, and Austria. Genome assemblies were constructed using PacBio HiFi and Hi-C sequencing data, yielding genome sizes of: 594.01 Mbp for *D. apicalis*; 587.80 Mbp for *D. pallida*; and 655.58 Mbp for *T. urticae.* Over 90% of sequences anchored into 13 pseudo-chromosomes per species. The assemblies for *D. apicalis* and *D. pallida* exhibited high completeness, capturing over 92% of conserved Hemiptera single-copy orthologues, as assessed by Benchmarking Universal Single-Copy Orthologues (BUSCO) analysis. Furthermore, we identified sequences of the primary psyllid symbiont, *Candidatus* Carsonella ruddii, in all three species. Comparative genomic analyses demonstrated synteny with other psyllid species. Notably, we observed significant expansions in gene families, particularly those linked to potential insecticide detoxification, within the *Dyspersa* lineage. Resequencing efforts also revealed the existence of multiple subpopulations of *D. apicalis* across Europe. These high-quality genome resources will support future research on genome evolution, insect-plant-pest interactions, and strategies for disease management.

**Significance:** Psyllid species are significant agricultural pests, known for transmitting plant diseases like *Candidatus* Liberibacter solanacearum (CLso), which causes ‘zebra chip’ in potatoes and ‘carrot yellows.’ However, genomic data on psyllids are limited. In this study, we present high-quality, chromosome-level genome assemblies for three psyllid species: *Dyspersa apicalis*, *Dyspersa pallida*, and *Trioza urticae*. We generated genome assemblies with over 90% of sequences anchored to 13 pseudo-chromosomes. Comparative analyses revealed gene expansions, particularly in detoxification pathways, suggesting adaptations within the Dyspersa lineage. Population resequencing of *D. apicalis* across Europe uncovered genetic subpopulations. These genomes will advance understanding of psyllid biology and inform disease management strategies.

## Introduction

Species from the insect superfamily Psylloidea (psyllids) are significant vectors of plant diseases in agriculture, transmitting pathogens that cause major economic losses. Psyllids are the exclusive vectors for *Candidatus* Liberibacter bacteria, including the infamous *Candidatus* Liberibacter asiaticus, which causes Huanglongbing disease in citrus (Huang et al. 2020). Other psyllid species, such as *Dyspersa pallida* (*Trioza anthrisci*), *Dyspersa apicalis* (*Trioza apicalis*), and *Trioza urticae* are known to transmit *Candidatus* Liberibacter solanacearum (CLso), a pathogen that affects solanaceous and apiaceous crops, leading to diseases such as ‘zebra chip’ in potatoes and ‘carrot yellows’ in carrots (Mishra and Ghanim 2022; Sjölund et al. 2017). *D. pallida* and particularly *D. apicalis* are a particular concern for carrot growers in Northen Europe as they are known to feed from carrot plants (Nissinen et al. 2021; Nissinen et al. 2022), whilst *T. urticae* has been found to transmit Lso haplotype U to nettle, but it’s impact on commercial crops appears to be negligible (Sumner-Kalkun et al. 2020). Interactions between plants, insects, and bacteria within the *Liberibacter* pathosystem are complex; different CLso haplotypes are tightly associated with specific psyllid hosts and their selectivity for particular plant hosts (Wang et al. 2017). Specific *Liberibacter* haplotypes and psyllid species may act symbiotically to suppress plant defence responses (Casteel et al. 2012). To date, fifteen CLso haplotypes have been identified; however, research on psyllid-associated microbes, including CLso, has primarily focused on phylogenetic barcoding studies, leaving much to explore regarding their ecological and evolutionary dynamics (Sumner-Kalkun et al. 2020a).

Psyllids, like many other Hemiptera, feed on plant phloem sap using their piercing-sucking mouthparts, and they may also feed on xylem sap when suitable phloem sources are unavailable (George et al. 2017). A recurring feature among sap-feeding insects, including psyllids, is their coevolution with bacterial endosymbionts, which play a crucial role in supplementing their nutrient-deficient diets (Moran and Bennett 2014). Although phloem sap is carbohydrate-rich it lacks essential amino acids, necessitating a reliance on endosymbiotic bacteria to synthesize these nutrients. These symbionts are transmitted vertically from mother to offspring, maintaining their evolutionary isolation. The primary symbiont of psyllids, *Candidatus* Carsonella ruddii (Gammaproteobacteria: Oceanospirillales), is believed to have maintained a mutualistic relationship with psyllids since their divergence from whiteflies over 240 million years ago (Spaulding and Von Dohlen 1998). *Ca*. C. ruddii has a highly streamlined genome and is entirely dependent on its hosts, residing in specialized bacteriocyte cells and relying on host support genes for survival (Moran and Bennett 2014). It represents an extreme case of genome reduction, with a genome size of just 160 kb, 182 open reading frames (ORFs), and a guanine-cytosine (GC) content of 16.5%, challenging the distinction between a cellular organism and an organelle (Nakabachi et al. 2006). Psyllids benefit from integrated essential amino acid biosynthesis pathways, combining genes from both the host and symbiont (Sloan et al. 2014). Horizontal gene transfer from bacteria to psyllid genomes is believed to have contributed to the extensive reduction in the Carsonella genome, which lacks functional genes for about half of the essential amino acid biosynthesis pathways (Sloan et al. 2014). Nearly all psyllids also harbour secondary symbionts, which may be facultative or obligate, but their identities can vary widely even among closely related taxa (Kwak et al. 2023).

Psyllids are vectors for *Ca*. Liberibacter and *Ca*. Phytoplasma bacteria, which are plant pathogens posing significant economic threats. Studies suggest that these pathogens can either enhance or diminish psyllid fitness, influenced by factors such as the nutritional quality of host plants and the degree to which these bacteria manipulate host plant processes to favour both themselves and their insect vectors (Mishra and Ghanim 2022). Different CLso haplotypes have been shown to have varying effects on psyllid fitness (Mishra and Ghanim 2022). The ability of different psyllid species to support various CLso haplotypes significantly impacts the risk of wild plants acting as disease reservoirs and the potential for spread of haplotypes to new climatic regions (Mishra and Ghanim 2022; Nissinen et al. 2022). *Ca*. Liberibacter and *Ca*. Phytoplasma bacteria must migrate from the alimentary canal to the salivary glands and multiply within the psyllid host to reach a sufficient threshold titre for transmission, consequently genetic diversity among psyllids is believed to impact the efficiency of pathogen transmission (Weil et al. 2020). Like *Ca*. C. ruddii, CLso has lost numerous genes related to metabolic and regulatory functions, and although this gene loss is less pronounced, neither *Ca*. C. ruddii nor CLso can be cultured in artificial media (Lin et al. 2011). Understanding the adaptations that allow these bacteria to colonize psyllids is essential for developing strategies to manage associated agriculturally important diseases. Moreover, insights into how *Ca*. Phytoplasma bacteria interact with their insect vectors have helped identify plant resistance genes, providing valuable knowledge for managing these pathogens (Huang et al. 2021).

The interaction between sap-feeding Hemiptera, such as psyllids, and their host plants involves the secretion of effector proteins in insect saliva, which act as virulence factors interacting with the plant immune system (Hogenhout and Bos 2011). There is a significant gap in our understanding of psyllid effector proteins which, to date, have only been studied in the Asian citrus psyllid, *Diaphorina citri* (Pacheco et al. 2020). Psyllids are thought to manipulate host plants through the secretion of these salivary effector proteins, and individual psyllid species often have specific associations with particular tissues of select host plants (Ouvrard et al. 2015). There are limited genomic resources available for studying psyllids, with genomes published for only two species of free-living psyllid, *D. citri* (Lei et al. 2024) and *Bactericera cockerelli* (Kwak et al. 2023), and one gall-forming psyllid, *Pachypsylla venusta* (Li et al. 2020), compared to approximately 53 aphid genomes available for the Aphidomorpha clade. Expanding these genomic resources is crucial for advancing our understanding of psyllid biology and their interactions with both plants and pathogens.

In this study, we have produced chromosome-level genome assemblies for *D. apicalis*, *D. pallida*, and *T. urticae*, effectively doubling the number of psyllid species with available genome sequences. Each of these species is known to vector CLso, with *D. apicalis* feeding on carrots and vectoring CLso haplotype C, *D. pallida* feeding on both carrots and wild plants, including cow parsley, also vectoring CLso haplotype C, and *T. urticae* feeding on nettles and vectoring CLso haplotype U. All three species are believed to overwinter in conifer trees (Kristoffersen and Anderbrant, 2007). Alongside the genome of the potato psyllid *B. cockerelli*, which vectors CLso haplotypes A and B, these new assemblies will facilitate research into the host specificity of CLso, as well as proteomic and transcriptomic studies to identify potential psyllid effectors that modulate plant processes and impact the efficiency of CLso transmission. This work significantly enhances the available genomic resources for psyllid research and opens new avenues for exploring the molecular interactions between psyllids, their symbionts, and host plants, ultimately contributing to the development of more effective disease management strategies.

## Materials and Methods

### Sample Collection

*D. apicalis* samples for *de novo* sequencing were collected from *Daucus carota* plants in Humppila, Finland, 2019 (∼ 60° 55’ 27.12”N, 23° 22’ 15.96”E) and were maintained in insectary colonies at Natural Resources Institute Finland (LUKE) in Jokionen until May 2021-temperature set to 20/15°C day/night, and photoperiod to 20:4 L:D (Nissinen et al. 2007), clean carrot plants cv. Fontana also grown in the same conditions – and then until December 2021 at JIC, Norwich, UK. We note that colonies of *D. apicalis* received from Finland could not be established at JIC until day length was increased from 16 hours to 18 hours. *D. pallida* samples were collected from *D. carota* plants in Elgin, UK, October, 2021 (∼ 57° 37’ 32.42737”N, - 3° 20’ 55.75437”E) and maintained as a colony on *Anthriscus sylvestris* at SASA, Edinburgh, UK until April 2022. Samples of *T. urticae* were collected from nettle in Norwich, UK, June 2021 and maintained in an insectary colony on nettle plants at JIC, Norwich, UK until October 2021. Psyllid individuals were retrieved from insectary cultures and frozen at −80° C prior to DNA extraction.

For resequencing, 40 *D. apicalis* and 9 *D. pallida* were collected (Supplementary Table S1, Supplementary Material online). Scottish *D. pallida* and Finnish *D. apicalis* samples were collected as described above. *D. apicalis* samples were also collected directly from field *D. carota* plants: in Thaur near Innsbruck, Austria, June 2021 (∼ 47° 17’ 41.14’’ N, 11° 27’ 40.81’’ E); and Larvik, Norway, August 2023 (∼ 59°7’16.8”N 10°3’40.9”E), in silica.

### DNA Extraction and Sequencing

For genome assembly, high molecular weight DNA was extracted from single individual psyllids using the Illustra Phytopure kit (GE Healthcare) as described by Mugford et al. 2020. A final DNA purification using AMPure beads (Beckman) according to the manufacturers protocol was used in place of isopropanol precipitation. DNA concentration was determined using Qubit DNA HS (Thermo-Fisher), purity by Nanodrop (Thermo-Fisher), and integrity by Femto Pulse (Agilent, performed by UCL Long-Read Sequencing Facility).

Tell-Seq (Chen et al. 2020) libraries were constructed using the Universal Sequencing Tell-Seq library preparation kit, using 5 ng input DNA according to the manufacturer’s instructions with minor modifications: 1 μl of tagging enzyme rather than 2 μl was used, and library cleanup was performed using 0.6-volumes of SPRI beads (Beckman) to optimise library insert size. Library titre was determined using the KAPA library quantification kit (Roche) using a CFX96 qPCR machine (Biorad), and library size was determined using a Tapestation High Sensitivity DNA kit (Agilent). Two libraries were pooled and sequenced on a NextSeq 550 using High Output Kit v2.5 (300 Cycles: 146 read 1, 18 index 1, 8 index 2, 146 read 2) (Illumina) according to the manufacturers protocol, with custom primers from the Tell-Seq kit.

The same DNA samples as used for Tell-Seq were sequenced by PacBio HiFi at the Norwegian Sequencing Centre. Libraries were prepared using the PacBio low input library kit (for *D. pallida* and *D. apicalis*) or the Ultra-low input kit (for *T. urticae*). The three libraries were pooled and sequenced across three 8M SMRT Cells.

For HiC sequencing, pooled samples of ∼30 (*T. urticae* and *D. apicalis*) or ∼10 (*D. pallida*) individuals -- from the same populations as the sample used for Tell-Seq and PacBio - were finely ground in a Retsch-mill with a steel ball bearing and resuspended in 4ml 1% formaldehyde for 20 minutes before quenching with the addition of 166 μl of 3M glycine, and incubated for 15 minutes. Samples were centrifuged at maximum speed for 1 minute, and the pellet was frozen. Proximo-HiC library preparation and sequencing were performed by Phase Genomics.

For resequencing, DNA was extracted from single psyllids as described in Sumner-Kulkun et al. 2020b, concentration and purity were determined by Qubit and Nanodrop (Thermo-Fisher). Samples were sequenced using Illumina 150PE reads, 8G per sample, by Novogene.

### Genome Assembly and Evaluation

PacBio HiFi reads were screened for internal adapters using HIFIADAPTERFILT (Sim et al. 2022). Following this, *de novo* assemblies were generated from Pacbio-HiFi data using HIFIASM v19.5.2 (Cheng et al. 2021) (parameters: “-l 1 --hg-size 820m --hom-cov 48 -D 10.0 -N 200 -s 0.25” for *D. pallida*; “-l 3 --hg-size 880m --hom-cov 29 -D 3.0 -N 200 -s 0.75” for *D. apicalis*; “-l 2 --hg-size 715m --hom-cov 12 -D 3.0 -N 200 -s 0.5” for *T. urticae*).

Tell-Seq reads were converted to 10X format using the ust10x tool provided by Universal Sequencing Technologies, internal barcodes and adapters were then trimmed using LONGRANGER v2.2.2 (https://github.com/10XGenomics/longranger). Reads were mapped to the draft assemblies with BWA-MEM v0.7.17 (Li 2013). The break10x command from SCAFF10X v4.2 (https://github.com/wtsi-hpag/Scaff10X) was used to split miss-joined scaffolds. Subsequently, trimmed Tell-Seq alignments were used to purge assemblies of haplotigs using PURGE_DUPS v1.2.5 (two rounds of chaining) (Guan et al. 2020) and PURGE HAPLOTIGS v1.1.2 with default settings (Roach et al 2018). K-mer spectra were plotted with MERQURY v1.3 (Rhie et al 2020) before and after purging of haplotig duplicates.

Hi-C reads were mapped to scaffolds via BWA-MEM v0.7.17 (parameters “-5SP-T0”) (Li 2013). High quality valid HiC pairs were identified (pairtools parse parameters “--min-mapq 40 --walks-policy 5unique --max-inter-align-gap 30”) and putative PCR duplicates removed (pairtools dedup) via PAIRTOOLS v0.3.0 (Open2C et al. 2023). YAHS v1.1 was used to scaffold the assemblies (Zhou et al. 2023). This resulted in chromosome level organisation for *D. pallida* and *D. apicalis* following manual curation using Pretext (https://github.com/wtsi-hpag/PretextMap; https://github.com/wtsi-hpag/PretextGraph; https://github.com/wtsi-hpag/PretextView). For *T. urticae* YAHS failed to scaffold the assembly successfully, scaffolds output from YAHS were therefore input to the 3DDNA scaffolding pipeline prior to manual curation with Pretext (Dudchenko et al. 2017). Finally, error correction and polishing for all assemblies was performed via Inspector (Chen et al. 2021).

### Mitochondrial Genome Extraction

Mitochondrial genome sequences were assembled using MITOHIFI v3.0 (Uliano-Silva et al. 2023). Mitochondrial sequences for *D. pallida* (NCBI: NC_038141.1) and *T. urticae* (NCBI: NC_038113.1) were used as reference sequences for their respective assemblies, with MitoHiFi running MitoFinder (Allio et al. 2020). For *D. apicalis*, since no pre-existing mitochondrial sequence was available, MitoHiFi was run with the parameter “-mitos” (Bernt et al. 2013) using *D. pallida* and *T. urticae* sequences as references. Both references highlighted the same contig as the *D. apicalis* mitochondrial genome.

### Non-Psyllid Sequence Identification

Sequences from the psyllid obligate symbiont *Ca.* C. ruddii were identified by low stringency BLASTN v2.9.0 search “E-value ≤ 1 × 10^-5^” of the *Ca.* C. ruddii genome (Genbank accession: GCA_000287275.1) against assembly scaffolds (Camacho et al. 2009). Scaffolds with hits were then investigated for collinearity with the *Ca.* C. ruddii genome via visualisation of MUMmer v4.0.0 alignments (Marçais et al. 2018).

Hi-C scaffolds were screened for contamination via the BLOBTOOLKIT v4.2.1 pipeline (Challis et al. 2020). Average coverage per scaffold was calculated by mapping trimmed Tell-Seq reads to the assemblies using BWA-MEM v0.7.17 (Li 2013). Each scaffold was annotated: with taxonomy information based upon BLASTN v2.12.0 (Camacho et al. 2009) searches against the National Center for Biotechnology Information (NCBI) nucleotide database (nt, downloaded October 2nd, 2023) with the parameters “-outfmt ‘6 qseqid staxids bitscore std’ -max_target_seqs 10 -max_hsps 1”; and with classifications by TIARA v1.0.3 (Karlicki et al. 2022). Resulting BAM, annotation table, and tiara files, as well as full BUSCO tables (hemiptera_odb10 database) (Simão et al. 2015) for each scaffolded assembly were then passed to BlobToolKit for plotting of “BlobPlots” and “SnailPlots”. Additional contamination screening was performed by taxonomic classification of all scaffolds via KRAKEN v2.1.3 (Wood et al. 2019) versus a database inclusive of GenBank, RefSeq, TPA and PDB (as of May 2nd, 2023) (downloaded September 15^th^, 2023 from https://benlangmead.github.io/aws-indexes/k2). Scaffolds classified to phyla other than Arthropoda were removed from assemblies. For *D. apicalis,* scaffolds with <0.3 or >0.4 GC content and <40 or >110 times coverage were removed unless they could be affirmatively classified to the Arthropoda taxa by kraken without contradictory classification by either blast or tiara. The same process was performed for *D. pallida* scaffolds with <0.3 or >0.4 GC content and <40 or >70 times coverage, and for *T. urticae* scaffolds with <0.3 or >0.45 GC content and <50 or >500 times coverage. Finally, contamination free status of final assemblies was confirmed via the GenBank Foreign Contamination Screen tool suite (Astashyn et al. 2024).

### Repetitive Element Classification

Repeat sequences were identified *de novo* via REPEATMODELER v2.0.5 (Flynn et al. 2020), and REPEATMASKER v4.1.5 with the parameters “-s -xsmall -html -gff -xm –species hemiptera” (Tarailo-Graovac et al. 2009). In addition, repetitive element families were quantified and kimura distance and repeat category plots via EARLGREYTE v4.0.8, utilising the Dfam 3.7 open database of transposable elements and repetitive DNA families, with RepeatMasker search term “Sternorrhyncha” (Baril et al. 2024).

### RNA Extraction and Sequencing

Total RNA was extracted from psyllids using Trizol (Merck) and RNeasy purification kit (Qiagen) with on-column DNAse treatment. Concentration and purity were determined by Qubit and Nanodrop (Thermo-Fisher) and integrity by Tapestation (Agilent). Library preparation and sequencing was performed by Novogene.

### Gene Prediction

Quality- and adapter-trimmed RNA-seq reads were mapped to soft-masked genome assemblies via HISAT2 v2.1.0 with parameters “--dta --max-intronlen 500000” (Kim et al. 2015), alignment files were then sorted and indexed via SAMTOOLS v1.18 (Li et al. 2009). Resulting BAM files were input as evidence to the BRAKER1 pipeline for each assembly (Hoff et al. 2016). ‘Arthropoda’ protein sequences from OrthoDB v11 (Kuznetsov et al. 2023) were supplemented with protein predictions from the Hemiptera clade and input as evidence to the BRAKER2 pipeline for each assembly (Brůna et al. 2021). TSEBRA was then used to combine BRAKER1 and BRAKER2 gene models, evidence weights “P = 1, E = 10, C = 5, M = 1, Intron support = 0, Stasto support = 1, e_1 = 0.5, e_2 = 2, e_3 = 0.1, e_4 = 0.36” were selected as providing the most complete gene sets (Gabriel et al. 2021). Additionally, the deep learning tool HELIXER v0.3.2, with the premade “invertebrate_v0.3_m_0100” model, was used to predict genes from each assembly (Holst et al. 2023). A combined protein sequence dataset was then compiled from the outputs of TSEBRA, Helixer, and ‘Arthropoda’ protein sequences from OrthoDB v11 supplemented with protein predictions from Hemiptera clade species.

Protein and transcriptomic evidence were input to the BRAKER3 pipeline to produce final gene predictions for each genome assembly (Gabriel et al. 2023). The quality and completeness of predictions were estimated via BUSCO and OMArk analysis (Simão et al. 2015; Nevers et al. 2024). Predicted proteins were searched against the curated portion of the UniProt protein database (Boeckmann 2003) using BLASTP v2.9.0 “E-value ≤1 × 10^−100^”(Camacho et al. 2009). INTERPROSCAN v5.52 was used to assign Gene Ontology (GO) terms and to annotate domains (Jones et al. 2014).

We also estimated the completeness of our genome annotations by re-aligning our RNA-seq data using STAR v2.5.0 (Dobin et al. 2013) and assessing the mapping rates to coding regions specifically (Supplementary Table S2, Supplementary Material online).

### Horizontally Transferred Genes

To identify horizontally transferred genes, genes highlighted as horizontally transferred by Sloan et al. 2014 were BLASTX v2.9.0 searched “E-value ≤ 1^-5^” against the *de novo* predicted proteins (Camacho et al. 2009). Additionally, predicted proteins were BLAST searched against the NCBI nr database (downloaded July 3^rd^ 2024) using DIAMOND v0.9.29 “E-value ≤ 1^-10^” (Buchfink and Huson 2015) and taxonomic classification of hits inspected with MEGAN v6.17.0 (Bağcı et al. 2021). Proteins classified to bacteria were aligned in Geneious v2023.1.2 (https://www.geneious.com) to identify homologues occurring across species.

### Phylogeny and Comparative Genomics

BUSCO sequences (hemiptera_odb10) were collected from 134 Hemiptera and 14 non-Hemiptera insect species genomes, including the available psyllid assemblies (5 *D. citri*; 2 *P. venusta*; 1 *B. cockerelli*; 1 *D. apicalis*; 1 *D. pallida*; 1 *T. urticae*) (Simão et al. 2015). Orthologues were aligned via MAFFT v7.520 (Katoh and Standley 2013) and trimmed with TRIMAL v1.4.1 (Capella-Gutiérrez et al. 2009). The Hemiptera phylogenetic tree was then calculated using IQ-TREE v2.3.0 (Minh et al. 2020) with parameters “-m GTR+F+I+R10-B 1000”.

Genes were predicted from *Ca.* C. ruddii genome assemblies from different psyllid hosts, including those screened from the *D. apicalis, D. pallida,* and *T. urticae* assemblies, using PROKKA v1.14.6 (Seemann 2014). ORTHOFINDER v2.5.4 (Emms and Kelly 2015) was used to generate a concatenated alignment of common orthologs. The *Ca.* C. ruddii phylogenetic tree was then calculated using IQ-TREE v2.3.0 (Minh et al. 2020) with parameters “-m mtInv+F+I+R4-B 1000”.

Orthology between Hemiptera protein predictions was assessed via ORTHOFINDER v2.5.4 (Emms and Kelly 2015). The longest isoform per gene for six Aphidomorpha, Heteroptera and Psylloidea species, along with representatives across the Hemiptera clade, and the hemipteroid *Frankliniella occidentalis*, were considered (Supplementary Table S3, Supplementary Material online). The Hemiptera phylogeny described above was pruned to contain only representatives of these 23 species and made ultrametric using R8S v1.7 (Mendes et al. 2021). Time points from the TimeTree database (http://timetree.org) were used for calibration of divergence time: *Myzus persicae-F. occidentalis,* 206-404.6 million years ago (MYA); *M. persicae-Aphis glycines,* 23-64.5 MYA; *M. persicae-Nilparvata lugens,* 112.5-391.7 MYA; and *M. persicae-Planococcus citri,* 168-286.4 MYA. Computational Analysis of gene Family Evolution (CAFE) v5.1 was used to characterise expansions and contractions of orthologous gene families (Mendes et al. 2021). Gene families with more than 100 members in any species were discarded prior to expansion/contraction analysis and an error model was estimated prior to estimation of a final birth-death (λ) parameter. Significantly expanded and contracted gene families in the *Dyspersa* clade were annotated via eggnog-mapper (Huerta-Cepas et al. 2019) and enrichment of KEGG terms was assessed using the R package “clusterProfiler” v4.10.1 (settings: pvalueCutoff = 0.05, pAdjustMethod = “BH”, qvalueCutoff = 0.05, minGSSize = 10) (Yu et al. 2012).

### Synteny Analysis

Syntenic blocks of genes in the chromosomes of *D. apicalis, D. pallida, P. venusta, D. citri* and *B. cockerelli* were identified. All-against-all BLASTP searches were performed between and within annotated protein sequences from each assembly with the parameters “E-value ≤ 1^-10^ -num_alignments 6 -outfmt 6”, resulting blast output files were concatenated and input to MCSCANX v1.1 with default settings (Wang et al. 2012). Syntenic alignments were visualised in SynVisio (Bandi and Gutwin 2020).

### Resequencing

Short read Illumina sequencing data was generated for 40 *D. apicalis* and 9 *D. pallida* samples (Supplementary Table S1, Supplementary Material online). Raw sequencing reads were trimmed and adapters removed using TRIMGALORE v0.6.10 (Krueger et al. 2023). BWA-MEM v0.7.17 was used to align trimmed reads to the *de novo D. apicalis* assembly and to the genome of *Ca.* C. ruddii (Accession no.: GCA_002009355.1) (Li 2013). PICARD TOOLS v2.1.1 was used to prepare sequence dictionaries for the two reference genomes and to mark and remove duplicate reads from alignment files, GATK v3.8.0 was used to realign reads near to indels and remove alignment artifacts (McKenna et al. 2010). Variants were called via SAMTOOLS/BCFTOOLS v1.15.1 “mpileup” and “call” commands (Li et al. 2009; Danecek et al. 2021).

Variants were filtered via GENMAP v1.3.0 and VCFTOOLS (Danecek et al. 2021; Pockrandt et al. 2019). Regions of the *de novo D. apicalis* that were repetitive (see section 2.6) or had below maximum mappability (GenMap with parameters “-K 100 -E2”) were masked (Pockrandt et al. 2019), only variants within unmasked regions were considered further. *D. apicalis* variants were filtered with VCFTools with parameters “--remove-indels -- max-alleles 2 --mac 1 --max-missing 0.9 --minQ 30 --minDP 5 --min-meanDP 5 --maxDP 40 --max-meanDP 40” (Danecek et al. 2021). *Ca.* C. ruddii variants were filtered with parameters “--remove-indels --max-alleles 2 --mac 1 --max-missing 0.9 --minQ 30 --minDP 5 --min-meanDP 5 --maxDP 300 --max-meanDP 300” (Danecek et al. 2021). Remaining biallelic SNPs were considered to be high confidence.

P-distance matrices were calculated from variant call format files via VCF2DIS v1.47, neighbour-net networks were then constructed using SPLITSTREE v4.14.8 (He 2022). Weir and Cockerham weighted F_ST_ was calculated between *D. apicalis* and *D. pallida* samples using VCFTools (Danecek et al. 2021). dXY between *D. apicalis* and *D. pallida* samples was calculated using a custom python script (https://github.com/TCHeaven/Scripts/tree/main/NBI/ calculate_dxy.py).

## Results

### Chromosome-scale Assembly of *D. apicalis, D. pallida,* and *T. urticae*

Chromosome level genome assemblies have been generated for *D. apicalis, D. pallida* and *T. urticae* utilizing PacBio HiFi and Tell-Seq data derived from single individuals of each species, as well as Hi-C data to facilitate higher level scaffolding. PacBio sequencing of *D. apicalis,* resulted in 2,847,503 HiFi reads, corresponding to a genome coverage of approximately 40×, additionally 189,033,534 Hi-C and 375,217,648 Tell-Seq reads were generated (Table 1). For *D. pallida* 2,454,894 HiFi reads were generated, equating to 32× coverage, along with 252,795,080 Hi-C reads and 438,125,030 Tell-Seq reads, whilst sequencing coverage of 35× was achieved for *T. urticae* with 3,052,430 HiFi reads along with 257,724,356 Hi-C reads and 793,362,872 Tell-Seq reads.

**Table 1.**
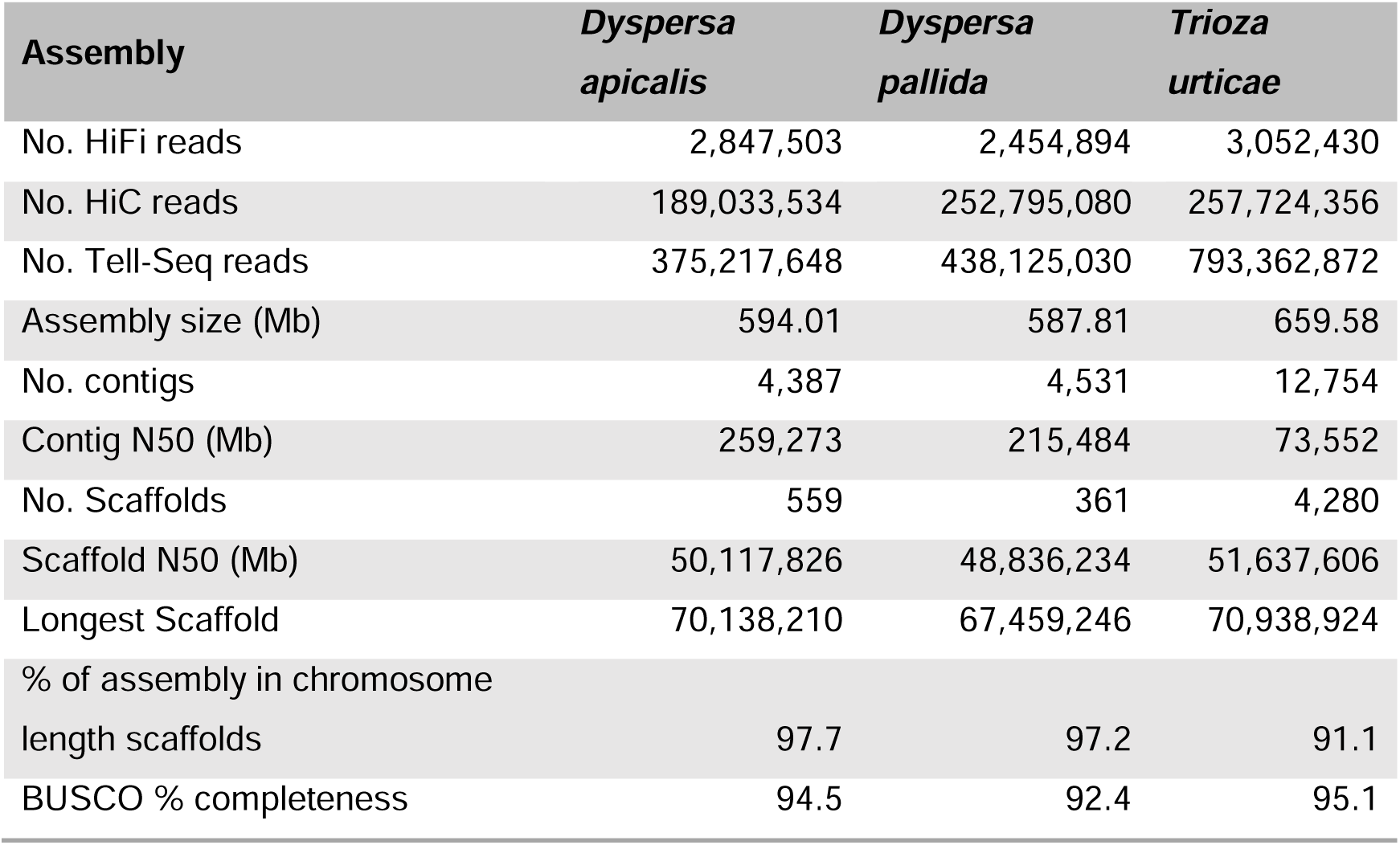
Sequencing and Assembly Metrics for Each Genome. . The table lists: the number PacBio HiFi, Tell-Seq, and HiC reads; final assembly size; number of sequences, and N50 before and after HiC scaffolding and curation; longest scaffold; percentage of assemblies anchored into pseudo-chromosome sequences; and the percentage of Hemiptera Benchmarking Universal Single Copy Orthologues present.

We generated chromosome length super scaffolds of the *D. apicalis* genome (Fig. 1A). This was achieved by an initial *de novo* assembly of *D. apicalis* HiFi reads with HiFiasm resulted in 4,387 scaffolds with an N50 of 259,273 bp. Following the removal of haplotigs, this assembly contained minimal duplication (3.9% of BUSCOs) (Fig. 1E) and was further scaffolded using Hi-C reads to generate chromosome length super scaffolds (Fig. 1A). K-mer analysis confirmed that purging minimised duplication without sacrificing unique sequence (Fig. 1E). After manual curation, and screening for symbiont and other non-psyllid sequences, the final *D. apicalis* assembly comprised 594.23 Mbp, with >97% of the assembly anchored into 13 chromosomes. Chromosomal scaffolds ranged from 25.9 Mbp to 70.1 Mbp in size. A total of 71 scaffolds were removed from the assembly during screening, including the mitochondrial genome.

**Fig.1.**
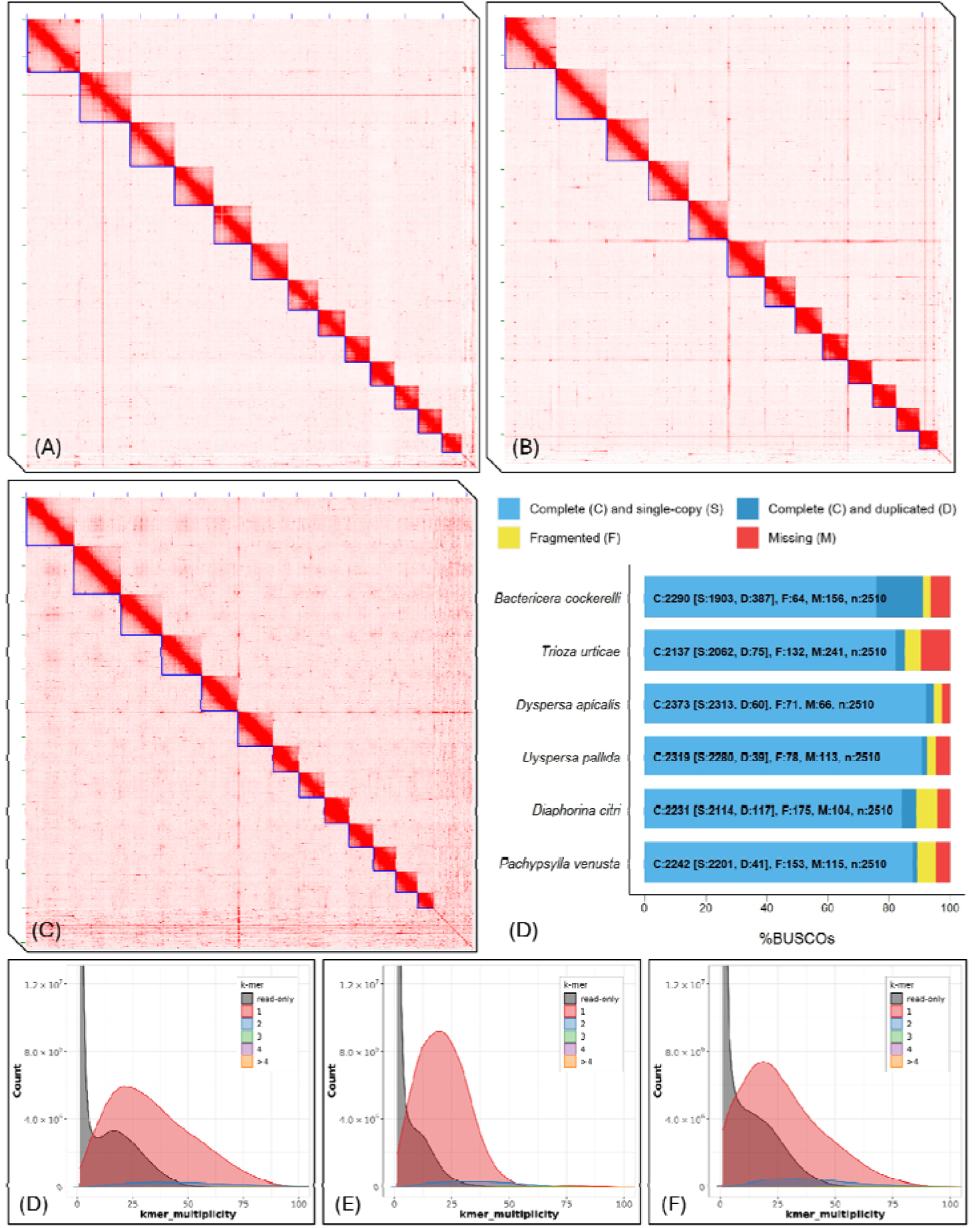
Chromosome level assemblies of *Dyspersa apicalis, Dyspersa pallida* and *Trioza urticae* were generated. Hi-C contact maps of *D. apicalis* **(A)***, D. pallida* **(B)** and *T. urticae* **(C)** display 13 super-scaffolds, indicated by blue lines, corresponding to 13 psyllid chromosomes. When assessed by Benchmarking Universal Single Copy Ortholog (BUSCO) analysis **(D)**, these assemblies have high completeness and low duplication compared to published psyllids genomes. *K*-mer spectra plots support minimal duplication of assemblies: *D. apicalis* **(E)***, D. pallida* **(F)** and *T. urticae* **(G)**. Gray shaded areas represent *k*-mers present in sequencing reads but not an assembly whilst red shaded areas represent *k*-mers found once in an assembly, other colours represent *k*-mers found multiple times in an assembly.

We found that the genome of *Ca.* C. ruddii was mistakenly integrated into the genome assembly of *D. apicalis*. To assess the boundaries of this misassembly, HiFi reads were aligned to the final *D. apicalis* assembly via MINIMAP v2.24-r1122 (Li 2018) and subsequent alignments visualised via QUALIMAP v2.2.2 and IGV v2.16.1 (Okonechnikov et al. 2016). This revealed that an approximately 170 kb region of the *D. apicalis* assembly had collinearity with the *Ca.* C. ruddii genome. Moreover, this region of the *D. apicalis* assembly also displayed markedly lower GC content and enormously higher sequencing coverage (∼1000× versus 40×) than surrounding regions. An inspection of read alignments also revealed that the region was flanked by a scaffold break and a position with ∼1× coverage only (Supplementary Figure S1, Supplementary Material online). The 170 kb region corresponding to the *Ca.* C. ruddii sequence was therefore excised from the scaffold.

Evaluation via Benchmarking Single Copy Orthologues (BUSCOs) confirmed that the *D. apicalis* assembly was highly complete with 94.5% of Hemiptera orthologues represented (Simão et al. 2015) (Fig. 1D). Indeed, this is the highest number of complete and single copy BUSCOs of any psyllid genome published to date.

Similarly, a chromosome-level assembly of *D. pallida* was generated (Fig. 1B). Initial *de novo* assembly of *D. pallida* HiFi reads resulted in 4,531 scaffolds with an N50 of 215,484 bp, with low duplication following haplotig purging (2.3% BUSCOs) (Fig. 1F), these were scaffolded via Hi-C data and manually curated. Following removal of mitochondrial scaffolds, *Ca. C. ruddii,* and other suspected non-psyllid scaffolds, the final assembly was 587.80 Mbp in length, with > 97% anchored into 13 chromosomal scaffolds ranging from 24.7 Mbp to 67.4 Mbp in size (Fig. 1B).

Forty-one sequences were classified as *Rickettsia*-like during screening of the *D. pallida* assembly, these possibly represent secondary symbionts. However, the largest of these sequences was only 88,053 bp in length. No *Rickettsia*-like sequences were identified in either *D. apicalis* or *T. urticae.* The *D. pallida* assembly was highly complete with 92.4% of Hemiptera orthologues represented (Simão et al. 2015) (Fig. 1D).

Initial *de novo* assembly of *T. urticae* resulted in 12,754 scaffolds, N50: 73,552. In the case of *T. urticae,* BUSCO duplication remained high following haplotig purging with Purge_Dups, additional purging was therefore performed using Purge Haplotigs (Roach et al. 2018), reducing BUSCO duplication to 3.7%. *K-*mer analysis confirmed that the assembly had very low levels of duplicated content following two rounds of haplotig removal (Fig. 1G). Hi-C scaffolding and manual curation resulted in 4,884 scaffolds, 604 of which were removed as potential non-psyllid sequences, including a complete *Ca. C. ruddii* genome. *Ca*.

C. ruddii sequences from both *D. apicalis* and *T. urticae* were assembled to distinct scaffolds which were removed from the psyllid assemblies. Whilst the final *T. urticae* assembly constituted 659.58 Mbp with 91% anchored into 13 chromosome scale scaffolds (Fig. 1C), BUSCO analysis revealed that only 83.9% of Hemiptera orthologues were present (Fig. 1D). The *D. apicalis* and *D. pallida* genomes are thus of a similar size to closely related psyllids such as *B. cockerelli* (567 Mbp) whilst the *T. urticae* assembly is the largest psyllid genome sequenced to date despite being incomplete (Kwak et al. 2023).

### Assemblies Feature High Repetitive Element Content

Masking and categorization of repetitive elements within the three psyllid genomes revealed that approximately two-thirds of each consists of Transposable Element (TE) sequences (Fig. 2). This contrasts to approximately one-third in *D. citri*, one-half in *P. venusta* and *B. cockerelli*, and 24.4% [±12.5% SD] across Hemiptera, Hymenoptera, Coleoptera, Lepidoptera and Diptera (Supplementary Figure S2-S5, Supplementary Material online). This expansion of transposable elements may drive the increased size of the *Dyspersa* and *T. urticae* genomes versus previously sequences psyllids (266-567 Mbp) (Kwak et al. 2023; Li et al. 2020; Lei et al. 2024). It was previously reported that the genome of *B. cockerelli* has relatively high TE content (Kwak et al. 2023). We classified an even larger proportion of the *B. cockerelli* genome as TEs (Supplementary Figure S5, Supplementary Material online). Additionally, the proportion of SINE elements was markedly lower in our analysis. This discrepancy may be due to the high levels of redundancy in SINE models generated by RepeatModeler2 and EarlGreyTE’s improved ability to characterize TE families (Baril et al. 2024).

**Fig. 2.**
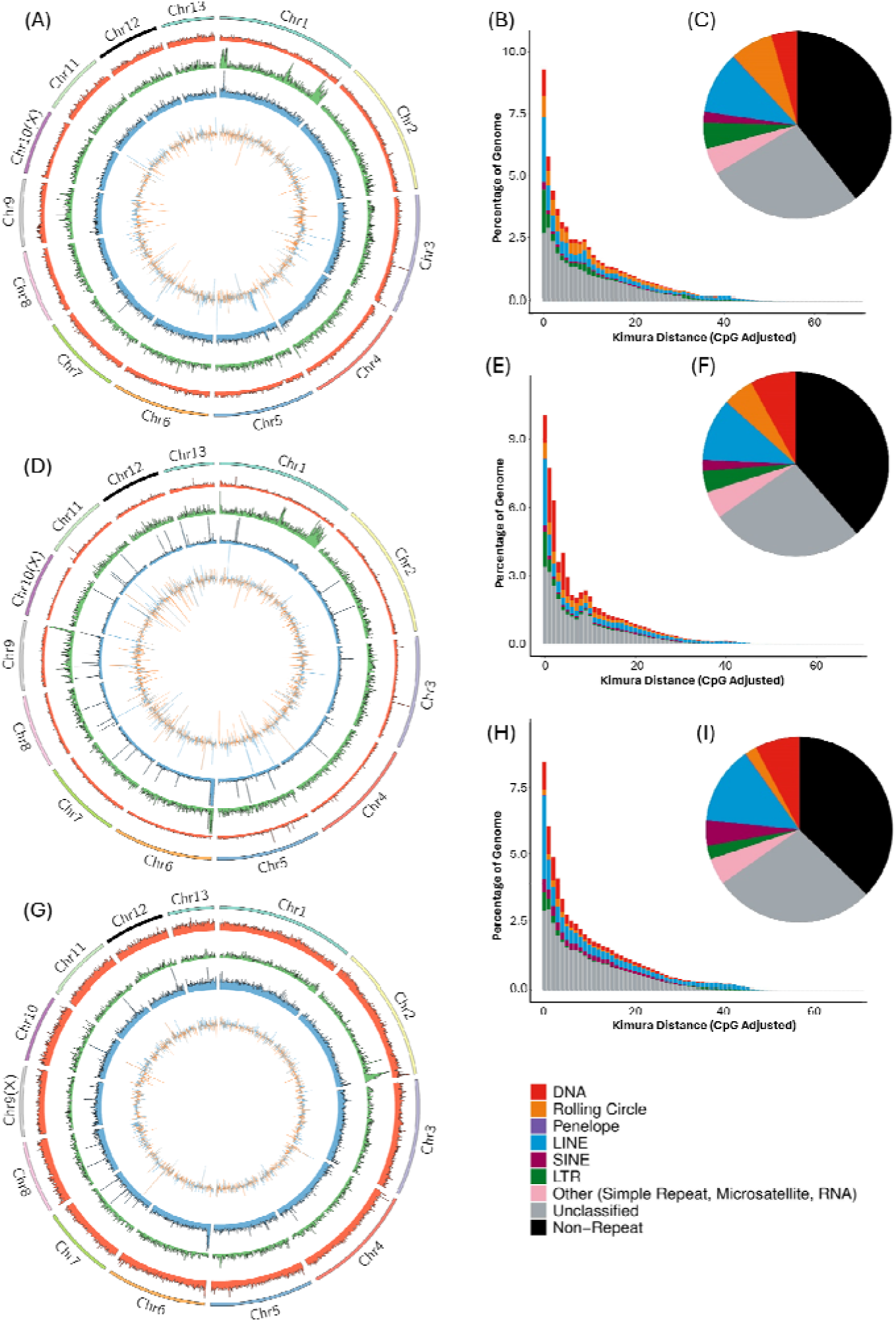
*Dyspersa* and *Trioza* psyllids have high Transposable Element (TE) content. Circos plots show the distribution of TEs, from outer to inner circles: 13 chromosome super-scaffolds; TE density per 200 Kb window; number of genes per 200 Kb; coverage represented by number of reads per 200 Kb; and guanine-cytosine (GC) content per 200 Kb are displayed for *Dyspersa apicalis* **(A)**, *Dyspersa pallida* **(D)**, and *Trioza urticae* **(G)** respectively. Kimura distance plots indicate the relative activity of different TE superfamilies in the genomes of *D. apicalis* **(B)***, D. pallida* **(E)**, and *T. urticae* **(H)**, more recently active elements are plotted towards the left hand side, whilst pie charts show the TE content of *D. apicalis* **(C)***, D. pallida* **(F)**, and *T. urticae* **(I)** assemblies, different colours represent different TE superfamilies.

Ancient transposon expansions are rare in the three psyllid genomes, with relatively recent transposon activity predominating (Fig. 2B,E,H). Notably, rolling circle elements are enriched, especially in the genomes of the two *Dyspersa* species. Our results are consistent with previous findings, which suggest that while TE content in insect genomes can vary significantly both between and within orders, it generally correlates with phylogenetic relatedness. This correlation is possibly due to the ability of TEs to transfer between closely related species. For example, at least one third of TEs in *Drosophila melanogaster, Drosophila simulans,* and *Drosophila yakuba* have been acquired through horizontal transfer (Bartolomé et al. 2009). These *Drosophila* species diverged approximately 11 million years ago, whereas *D. apicalis* and *D. pallida* may have separated less than 5 million years ago (based on the time calibration of our Hemiptera phylogeny). Therefore, and given the close evolutionary relationship and ecological crossover of *D. apicalis* and *D. pallida,* similarities in TE content are unsurprising.

### Detoxification Gene Families are Expanded

In total, 22,637 genes were predicted in the *D. apicalis* genome assembly, 22,592 in the *D. pallida* assembly and 19,331 in the *T. urticae* assembly (Table 2). This gene count is comparable to previously sequenced psyllids. For instance, *D. citri* has 19,083 to 24,730 annotated protein-coding gene models (Lei et al. 2024). Gene annotations were generated using the BRAKER3 annotation pipeline, incorporating predictions from BRAKER1, BRAKER2, and Helixer as protein evidence. This approach was found to yield the most complete annotation sets while minimizing unexpected gene duplication (Supplementary Table S4, Supplementary Material online). Analysis with both BUSCO and OMArk indicated that the proteomes for *D. apicalis* and *D. pallida* were highly complete, whereas *T. urticae* exhibited a higher level of missingness (Fig. 3A,B,C). This high level of completeness was achieved despite a relatively low mapping rate (∼60%) between RNA-Seq data and the genome assembly in each case. The percent of unmapped reads was higher for our *T. urticae* samples (25% of male reads; 27% of female reads) than for the other species (12-17% for *D. apicalis*; 19-23% for *D. pallida*; Supplementary Table S1, Supplementary Material online). The unmapped reads were too short to be properly aligned, indicating that this was due to library quality and not contamination or problems with our assemblies. In total, 11,059 *D. apicalis*, 10,639 *D. pallida*, and 8,921 *T. urticae* transcripts were annotated with at least one Gene Ontology (GO) term.

**Fig. 3.**
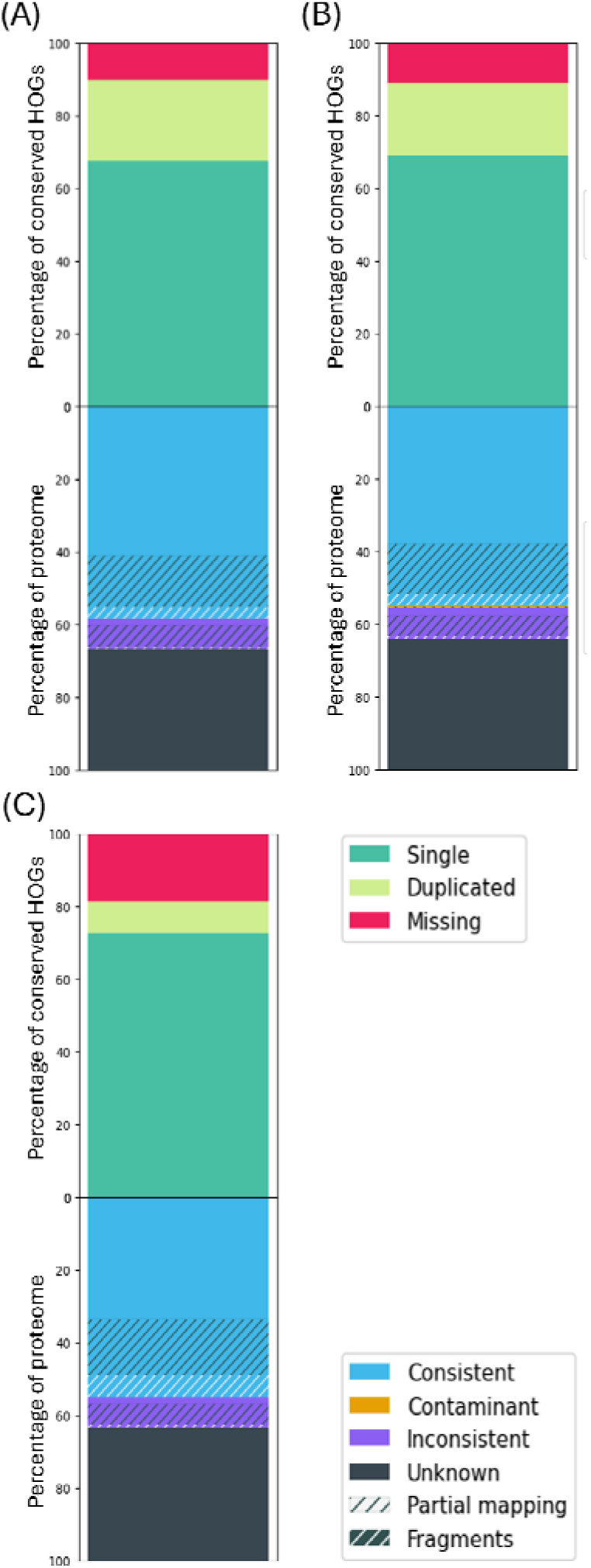
The annotated *Dyspersa* proteomes are highly complete. The completeness and consistency of Hierarchical Orthologous gene Groups (HOGs) for each assembly - *Dyspersa apicalis* (A), *Dyspersa pallida* (B), and *Trioza urticae* (C)—are plotted. Completeness is represented in the top stacked bar plot, showing the percentage of single copy (green), duplicated (yellow), and missing (red) genes. Consistency is represented in the bottom stacked bar plot, displaying the percentage of taxonomically consistent (blue) and inconsistent (purple) genes. Contaminant genes are shown in orange, while genes with no taxonomy within the OMArk database are in black.

**Table 2.**
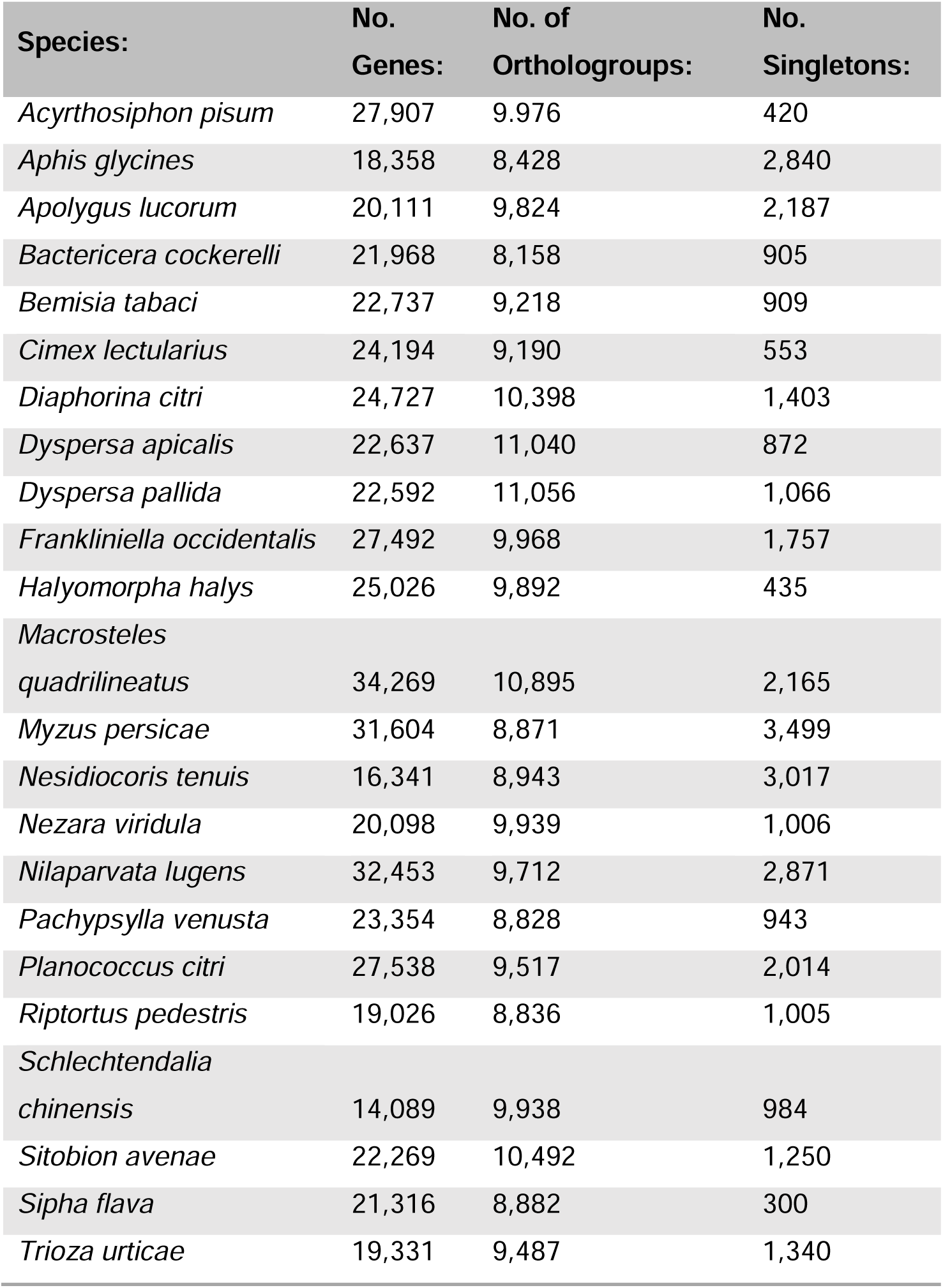
Gene content of Psyllid assemblies. The number of genes, orthogroups, and singletons predicted across Hemiptera species are given.

Phylogenetic analysis confirmed that *D. apicalis* and *D. pallida* are closely related species within the Psylloidea clade, with an estimated divergence time less than 5 million years ago (based upon R8S phylogenetic tree calibration) (Fig. 4). *T. urticae* was found to be most closely related to *B. cockerelli,* whilst *D. citri* and the gall forming psyllid *P. venusta* were shown to be more distantly relatives. Additionally, the phylogeny of *Ca.* C. ruddii symbionts from different psyllids is congruent with the phylogeny of their respective host insects, reflecting the maternal transmission of the symbiont (Supplementary Figure S6, Supplementary Material online). These results are consistent with previous studies (Percy et al. 2018).

**Fig. 4.**
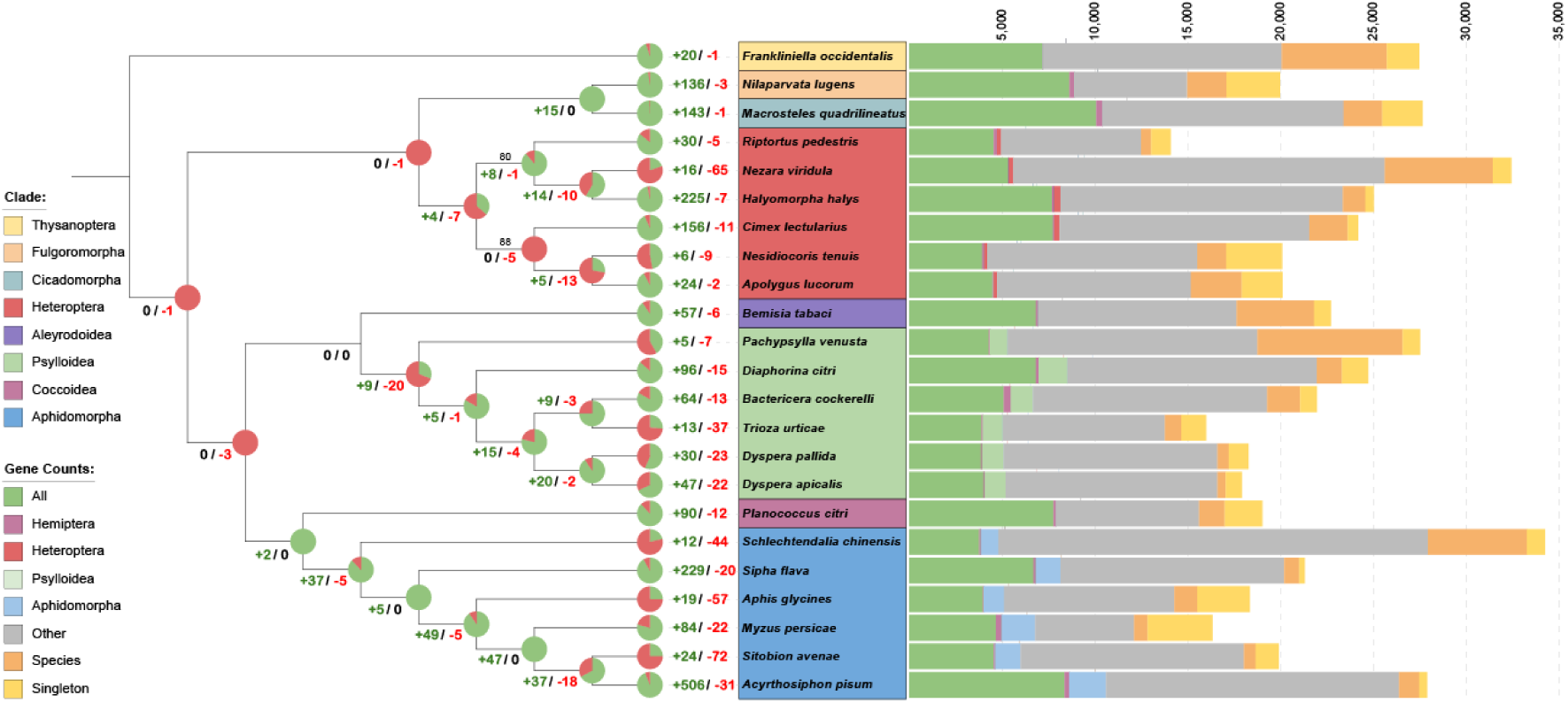
Psyllid lineages are characterised by expanding gene families. The orthology of protein predictions across seven Hemiptera infra-orders is considered. A stacked bar chart shows the number of genes for each species corresponding to orthogroups shared across: all included Hemiptera and *Frankliniella occidentalis*; all Hemiptera but not *F. occidentalis*; five out of six members of the phloem feeding infra-orders Psylloidea and Aphidomorpha, but not any non-phloem feeding species; five out of six members of the Heteroptera, Psylloidea, or Aphidomorpha infra-orders but absent from all other species; other combinations of different species; one species only; as well as singletons with no orthogroup placement. Pie charts display the number of orthogroups undergoing significant (p ≤ 0.01) expansion or contraction at different divisions of the Hemiptera phylogeny.

Orthology clustering was performed for the proteomes of *D. apicalis*, *D. pallida*, and *T. urticae* (the complete set of annotated protein-coding genes), along with proteomes of 19 other Hemiptera species and the thrips *F. occidentalis* (Supplementary Table S2, Supplementary Material online). The longest isoform for each gene was considered, resulting in clustering of 483,650 genes (93.5 % of all) into some 31,659 orthogroups. *D. apicalis, D. pallida,* and *T. urticae* proteins were allocated into 11,040, 11,056, and 9,487 orthogroups respectively (Table 1; Supplementary Table S5, Supplementary Material online). A total of 2,549 orthogroups were represented in all 23 species, constituting gene families fundamental to the Hemiptera clade. Orthologues specific to psyllids were investigated, revealing 859 orthogroups common across at least five out of six psyllid species sequenced to date but absent from other Hemiptera and *F. occidentalis* (Fig. 4). These orthogroups likely arose early following the divergence of the Psylloidea clade and probably include proteins essential for supporting the ubiquitous psyllid endosymbiont *Ca*. C. ruddii. The number of psyllid-specific orthogroups was comparable to the number aphid specific orthogroups (811). In contrast, heteropteran species, which do not share the phloem-feeding lifestyle of psyllids and aphids, only shared 162 infra-order specific orthogroups.

Gene family expansion and contraction analysis revealed 69 orthogroups to be significantly (p ≤ 0.01) expanded or contracted in the *D. apicalis* lineage whilst 53 orthogroups were expanded or contracted in *D. pallida* (Fig. 4; Supplementary Table S6, Supplementary Material online). Our results support previous observations that the evolution of free-living psyllid lineages is characterized more by the expansion of gene families than by their contraction (Kwak et al. 2023; Lei et al. 2024). Conversely, *P. venusta* exhibits more contracted than expanded gene families. The *T. urticae* proteome contained 13 expanded and 37 contracted orthogroups; however, given the lower BUSCO and Omark scores for this assembly it is likely the *T. urticae* proteome is incomplete.

Genes within *D. apicalis* expanded orthogroups were significantly enriched (p ≤ 0.05) for six KEGG orthology terms: K13442 (calpain-14), K08811 (testis-specific serine kinase), K11251 (histone H2A), K11252 (histone H2B), K11275 (histone H1/5), and K15001 (cytochrome P450 family 4). In *D. pallida*, expanded orthogroups were enriched for four terms: K07378 (neuroligin), K15743 (carboxylesterase 3/5), K06515 (choline transporter-like protein), and K00699 (glucuronosyltransferase). Additionally, orthogroups expanded within the *Dyspersa* clade were enriched for four KEGG orthology terms: K13443 (Niemann-Pick C2 protein), K08811 (testis-specific serine kinase), K19679 (intraflagellar transport protein 74), and K05033 (ATP-binding cassette subfamily C member 9) (Supplementary Table S7; Supplementary Figure S7-S9, Supplementary Material online).

### Horizontally Transferred Genes

Streamlining of symbiont genomes can result in gene loss and disrupted biosynthetic pathways. In the psyllids *P. venusta, D. citri,* and *B. cockerelli* Horizontal Transfer of bacterial Genes (HTGs) into the insect genome compensates for *Ca*. C. ruddii gene losses, biosynthetic pathways for essential amino acids in psyllid species integrate genes from both host and symbiont genomes. We anticipated that the same HTGs would be present in *D. apicalis, D. pallida,* and *T. urticae* genomes given that they share the primary symbiont *Ca*.

### C. ruddii

We observed HTG orthologs first identified in the transcriptome of the gall-forming psyllid *P. venusta* by Sloan et al. amongst *D. apicalis, D. pallida,* and *T. urticae* gene predictions (Sloan et al. 2014). These included ORF (AAA-ATPase-like), RSMJ (rRNA methyltransferase), ASL-1 (ArgininoSuccinate Lyase), ASL-2, CM (Chorismate Mutase), MUTY, and YDCJ (Sloan et al. 2014). *ASL* and *CM* genes are involved in the synthesis of the essential amino acids arginine and phenylalanine respectively. The acquisition of a bacterial CM gene is a crucial adaptation in psyllids, as whilst *Ca*. C. ruddii retains a *pheA* gene it has lost the CM domain (Sloan et al. 2014). Similarly, the *argH* gene encoded by *Ca*.

C. ruddii can no longer produce a functional ASL protein due to the loss of key catalytic residues (Tamames et al. 2007). In addition to previously described HTGs a gene encoding an Rpn family recombination-promoting nuclease/putative transposase with similarity to *Wolbachia* genes was predicted in both *D. apicalis* and *D. pallida*.

Neither RIBC (riboflavin synthase) nor Ankyrin repeat domain HGT proteins, which were identified in *P. venusta* (Sloan et al. 2014), were found in any of the three proteomes generated in this study. The ankyrin repeat domain gene appears to be confined to the *P. venusta* lineage, however we were surprised to find no *ribC* genes in the *Dyspersa* or *T. urticae* genomes as the expression of a *ribC* gene has previously been reported in *P. venusta*, *D. citri* and *B. cockerelli*, suggesting that its acquisition occurred early in the development of the psyllid clade (Sloan et al. 2014; Kwak et al. 2023). In *P. venusta*, the secondary symbiont *Candidatus* Profftella armatura carries all the genes required for riboflavin biosynthesis except *ribC*, enabling a complete pathway. However, *Ca.* P. armatura is not a known symbiont of *D. apicalis*, *D. pallida*, or *T. urticae*. It is possible that, in the absence of a suitable collaborating symbiont, the *ribC* gene has been lost from these lineages.

### Inter-Chromosomal Rearrangements are Rare in Psyllids

Assessment of gene synteny across available psyllid assemblies revealed a broadly conserved genome structure at the chromosome level (Fig. 5). Additionally, whilst intra-chromosomal rearrangement appears common between psyllid species from different genera, relatively few intra-chromosomal rearrangements were observed between *D. apicalis* and *D. pallida*. This likely reflects the recent divergence of the two species. The lack of inter-chromosomal rearrangements seen in psyllids contrasts with the situation in aphids where rearrangement is common (Mathers et al. 2021; Li et al. 2020). However, in both aphids and psyllids sex chromosomes are highly conserved (Mathers et al. 2021). Indeed, the sex chromosome (the 10^th^ largest chromosome) appears to have undergone no obvious rearrangements at all between *D. apicalis* and *D. pallida* (Fig. 5). Li et al. previously compared the X chromosomes of *Acyrthosiphon pisum*, *Rhopalosiphum maidis*, and *P. venusta* (Li et al. 2020). They reported striking differences between aphid and psyllid X chromosomes. Namely that, in aphids, X chromosomes are enriched for male-biased genes, whereas in psyllids, male-biased genes are enriched on autosomes. Aphids and psyllids share an X0 sex determination system. However, outside a few single-sex populations psyllids only reproduce sexually and, unlike many aphids, do not undergo parthenogenesis (Li et al. 2020). Specifically, *D. apicalis* and *D. pallida* are univoltine and reproduce once per year, whilst *T. urticae* is multivoltine in the UK (Davis 1973).

**Fig. 5.**
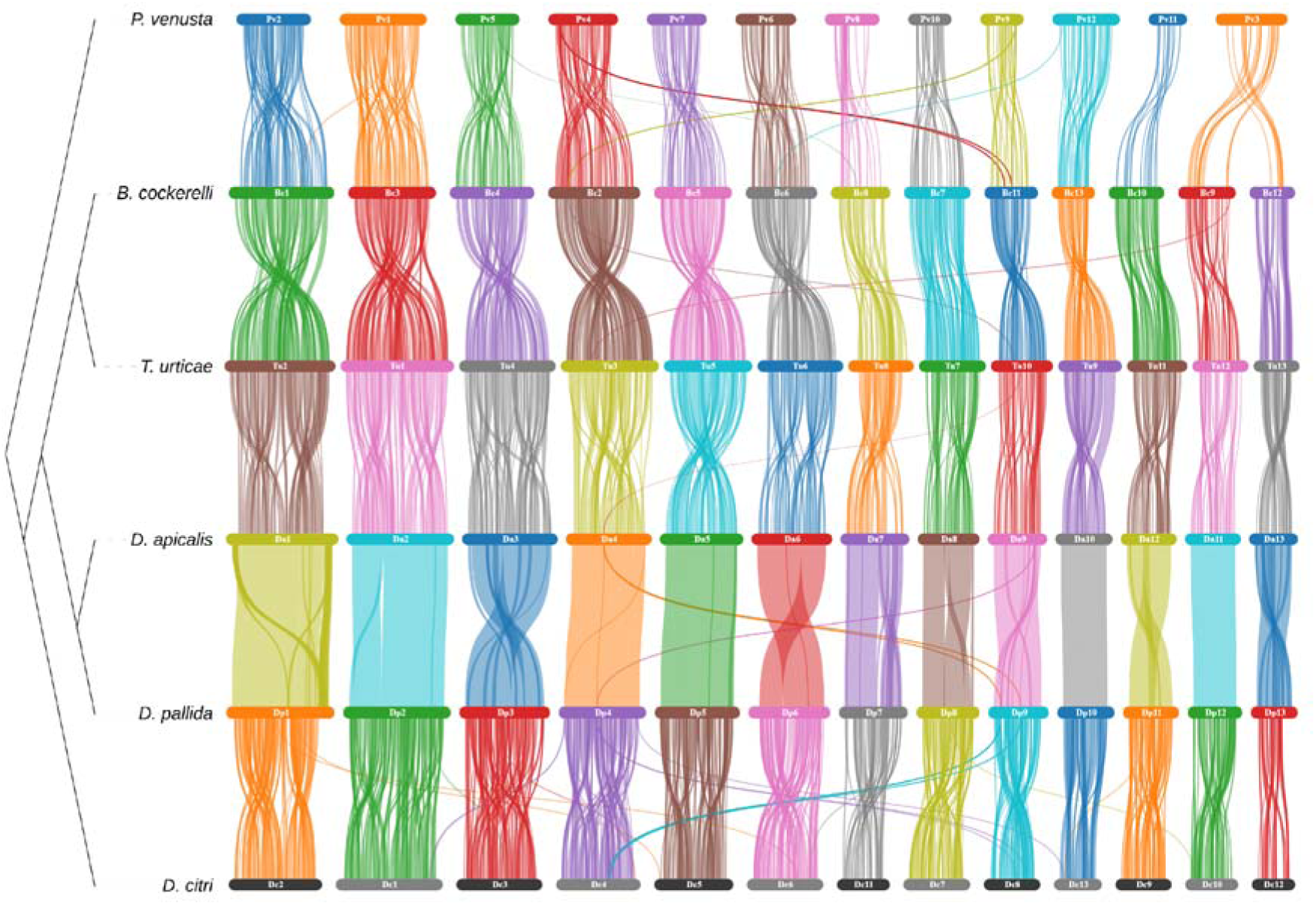
Chromosomal synteny between psyllid species. Species represented are, from top to bottom; *Pachypsylla venusta, Bactericera cockerelli, Trioza urticae, Dyspersa apicalis, Dyspersa pallida,* and *Diaphorina citri.* Connections are drawn between gene locations with reciprocal blast hits. Chromosomes are numbered in size order from largest to smallest and ordered in relation to *D. pallida*.

The *P. venusta* lineage forms a sister taxon to the other five sequenced psyllids and exhibits a markedly different lifestyle. Psyllids occur as gall-inducing, free-living, and lerp-forming taxa, as a gall forming species young *P. venusta* become completely encased within galls (Li et al. 2020). Consistent with previous reports, the free-living psyllids sequenced in this study have 13 chromosomes, whilst the gall forming psyllid *P. venusta* has only 12, constituting the only major inter-chromosomal rearrangement documented in psyllids to date (Li et al. 2020). Given the synteny between the chromosomes of free-living species and *P. venusta*, the difference in chromosome number is likely the result of either a chromosomal fission or fusion event. However, additional chromosomal-level psyllid assemblies from the *Pachypsylla* adjacent psyllid taxa will be required to determine the origins of this change.

### *Dyspersa apicalis* Population Structure

A total of nine *D. pallida* individuals from Scotland, as well as 41 *D. apicalis* individuals (12 Norwegian, 14 Finnish, and 15 Austrian) were sequenced. An average of 28,163,248 raw reads were generated for each individual, with an average genome coverage of 15.6× for *D. apicalis* and 11.7× for *D. pallida* samples. SNPs were called against the newly assembled *D. pallida* genome and reference *Ca.* C. ruddii assembly and were used to assess the population structure of the two psyllid species. A total of 9,674,852 high quality biallelic SNPs were called against the *D. pallida* genome and 9,226 against the *Ca.* C. ruddii genome.

SNPs called against both *D. pallida* and *Ca.* C. ruddii genomes group *D. pallida* separately from *D. apicalis,* indicating an extended period of reproductive isolation (Fig. 6A,B). This result supports the speciation of *D. apicalis* and *D. pallida*, as do F_ST_ (0.5625) and dXY (0.4745) measures of divergence which indicate high genomic differentiation. Congruence between the host and symbiont populations further supports vertical maternal transmission of the endosymbiont *Ca.* C. ruddii, as there is no evidence of insect-to-insect interspecies transmission despite the overlapping host and geographic range of the two species. Nonetheless, when compared to strains sampled from the wider psyllid clade *D. apicalis* and *D. pallida Ca.* C. ruddii endosymbionts are closely related (Supplementary Figure S6, Supplementary Material online).

**Fig. 6.**
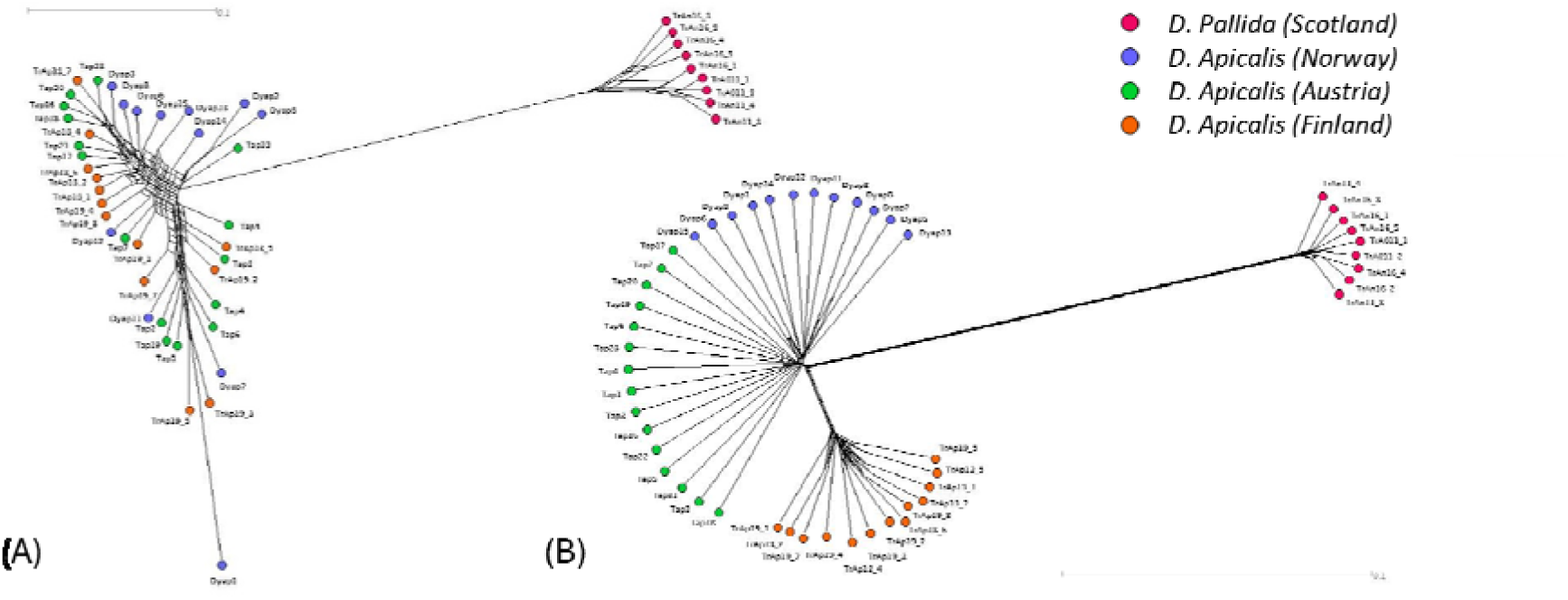
Genetic relatedness of European *Dyspersa apicalis* and *Dyspersa pallida* samples. Neighbour net networks are shown, these were calculated from single nucleotide polymorphisms called versus the *Candidatus* Carsonella ruddii genome **(A)** and *de novo D. apicalis* genome **(B).** *D. pallida* samples from Scotland (red) and *D. apicalis* samples from Norway (blue), Austria (green) and Finland (orange) are plotted.

Within *D. apicalis* samples, SNPs from *Ca.* C. ruddii present a different population structure compared to the SNPs from in the insect genome. Clustering of samples based upon SNPs in the psyllid genome results in separation of *D. apicalis* individuals from Finland which form a distinct population, whilst individuals from Norway and Austria are closely related (Fig. 6B). Conversely, *Ca.* C. ruddii SNPs suggest one large *D. apicalis* population (Fig. 6A). F_ST_ (0.15864) and dXY (0.0.2831) measures indicate moderate differentiation between the Finnish and Austrian/Norwegian *D. apicalis* populations. This separation of the Finnish samples is surprising given that environmental factors such as day length are more similar between sampling locations in Norway and Finland than between these Nordic countries and Austria. Peak summer solstice day lengths were 15h:56m:21s, 18h:32m:46s, 19h:14m:28s, and 18h:04m:05s respectively for the Austrian, Norwegian, Finnish, and UK sampling locations. Additionally, winds across Europe blow predominantly with an east-west, rather than north-south, direction and one might reasonably assume this would favour a longitudinal spread of winged insects over a latitudinal spread.

## Discussion

### Adaption in the *Dyspersa* genus

The genome features of psyllids influence their evolutionary adaptability, and the threat they pose as agricultural pests. We observe expansion of carboxylesterase and glucuronosyltransferase genes in *D. pallida,* and cytochrome P450 family 4 genes in the *D. apicalis*, this is notable because these gene families are known to facilitate insecticide detoxification (Yuan et al. 2021). Similarly, ATP-binding cassette subfamily C genes, which are expanded in the *Dyspersa* genus, have been associated with insecticide resistance ^128^. Insecticide resistance is a growing issue in *D. citri* due to the extensive use of pesticides to contain Huanglongbing disease (Wang et al. 2019). Analysis of gene evolution in *D. citri* also highlighted enrichment of cytochrome P450 genes amongst expanding gene families, presumably promoting the metabolism of toxic substances (Lei et al. 2024). It is plausible that similar adaptations are taking place in *Dyspersa* species. Suspected Pyrethroid resistance in *D. apicalis* has been reported in Norway, following several decades of heavy use (Nissinen et al. 2021). However, genes identified in studies of pesticide resistance may well have roles in the detoxification of natural plant defence compounds, and so the expansion of detoxification related gene families in *D. apicalis* and *D. pallida* may equally be adaptations to herbivory of their respective host plants. Either way, an increased number of detoxification genes provides more targets for potential adaptive mutations against insecticides in the future.

TE sequences are increasingly seen as catalysts of adaptive change and innovation in genomes (Gilbert and Feschotte 2018). We report that *D. apicalis, D. pallida,* and *T. urticae* genomes have comparatively high repeat content versus many other hemipteran species (Zidi et al. 2022). TE transfers between parasites and their hosts have been documented (Gilbert and Feschotte 2018). Exchange of genetic material between psyllids would reduce the probability that particular CLso haplotypes remain confined to individual psyllid species. Whilst hemipteran endosymbiont genomes are now highly streamlined it is tempting to speculate what role TE may have played in the historical acquisition of bacterial genes. TE-mediated adaptation can lead to the emergence of new phenotypes, as transposable elements often carry regulatory elements that can alter gene regulation (Gilbert and Feschotte 2018). In *Bemisia tabaci* it has been reported that TEs are inserted upstream of cathepsin genes (Zidi et al. 2022). The resources presented in this study will facilitate further research into the role of TEs in psyllid evolution.

### Finnish *Dyspersa apicalis* Sub*-*Population

An important question in the study of the CLso/psyllid pathosystem in Europe is why the severity of outbreaks varies between regions. For example, CLso outbreaks on carrot are a particularly serious problem in Finland but are not observed in the UK, despite the presence of *D. apicalis* in carrot growing areas. In the UK, CLso has so far only been found in asymptomatic carrot plants (Sumner-Kalkun et al. 2020) as well as parsley seeds (Monger and Jeffries 2016), and in the psyllid *D. pallida* rather than *D. apicalis* - the primary vector in Northern Europe, including Norway and Sweden (Sjölund et al. 2017). The easy availability of psyllid overwintering sites (70% of Finland is covered by forests) may result in more severe CLso outbreaks, or alternatively differences in CLso genetics between closely related haplotypes may be responsible (Nissinen et al. 2022). A further hypothesis is that different regional populations of psyllid may exist which vary in their competence as CLso vectors (Haapalainen et al. 2018). A latitudinal cline in the genetic diversity of *T. urticae* from Greece to Norway, with major haplogroups separated by natural geological barriers, has been described (Wonglersak et al. 2017). In contrast, *B. cockerelli* individuals in North America have been assigned to two large groups, one predominating in the west and one from the central USA to eastern Mexico (Swisher et al. 2014). We were therefore interested to investigate any differentiation between regional psyllid populations.

Our results appear to contradict a *D. apicalis* population structure based on latitude and imply the existence of multiple population groups but also the mixing/predominance of individual groups over large geographic areas. Finnish samples were observed to form a distinct population. It is possible that the isolation of Finnish *D. apicalis* individuals may represent a bottleneck during the period these samples were maintained as an insectary colony. Alternatively, we found that day lengths of over 16 hours are required to maintain Finnish originating *D. apicalis* lineages (Victor Soria-Carrasco, personal communication), a photoperiod which never occurs at the Austrian sampling site, implying some heritable adaptation to latitude must be present in Finnish samples as in Swedish population (Valterová et al. 1997). It is conceivable that a differentiated Finnish *D. apicalis* population is a better host for CLso, but is unable to survive at lower latitudes with shorter maximum day length.

Photoperiod is believed to be the main determinant affecting the production of different seasonal forms of psyllid, including the migration of *D. apicalis* from conifer trees to carrot in May/June (Kristoffersen and Anderbrant 2007). However, this interaction appears to break down at higher latitudes, potentially being mediated by temperature (Butterfield et al. 2001). Whilst the flight capacity of *D. apicalis* is not well known, a study of the epidemiology of CLso suggests that infected *D. apicalis* may travel 5-10 km from winter to summer hosts (Nissinen et al. 2022). Migration of insect vectors affects the epidemiology of diseases such as CLso. The life cycle of *D. apicalis* is analogous to that of *Cacopsylla pruni,* the vector of *Candidatus*

Phytoplasma prunorum. In the *C. pruni – Ca.* P. prunorum system bacterial strains are disseminated over long distances as *C. pruni* must migrate to conifer trees over winter and are only competent to transmit *Ca.* P. prunorum following an eight month incubation period (Thébaud et al. 2009). Similarly, *D. apicalis* overwinters on conifers and produces annual broods of offspring (Láska 2011). Although, in contrast to the *C. pruni* eight month incubation period, the latency period of CLso haplotypes A and B is reportedly only 17-25 days in *B. cockerelli* (Tang et al. 2020).

In this study samples were collected from only one field in each country considered. Notably, whilst CLso has caused significant yield reductions for carrot growers in Finland and Norway (Haapalainen et al. 2017), the only location where CLso is known to occur in Austria is the Inn valley near Innsbruck, Tyrol (Lethmayer and Gottsberger 2020). The source of the Austrian infection is unknown. Given that individuals of Austrian and Norwegian *D. apicalis* populations group together and separately from those of Finland (Fig. 6B), it is possible that an expatriate population of *D. apicalis* originating from Norway has established itself within the Inn valley. This would explain the emergence of CLso in the area and the relatedness of Norwegian and Austrian individuals. However, given the formidable natural barriers between Norway and Austria natural dispersal of a single *D. apicalis* haplotype between the two regions seems unlikely. The possibility of trade related dispersal should be considered. Further surveys with more broad-based sampling will provide greater clarity on the true population structure of *D. apicalis* across Europe.

## Conclusions

Psyllids are important pests of world agriculture due primarily to their ability to vector phytopathogenic bacteria. This study provides the first genomic resources for *D. apicalis* which is a particular problem for growers of apiaceous crops in Europe, as well as *T. urticae* and *D. pallida* affecting neighbouring wild plants. These assemblies will facilitate further research into the biology of *Ca.* liberibacter solanacearum, the cause of ‘zebra chip’ disease in potato and ‘carrot yellows’ disease in carrot. The high quality of the psyllid genome assemblies, and accompanying *Ca.* C. ruddii sequences, will also allow further investigation of genome evolution and symbiosis in Hemiptera. Additionally, we found that *D. apicalis* in Finland has adapted to long daylength conditions, and resequencing of *D. apicalis* samples from across Europe suggests that the Finnish populations groups separately from those of Norway and Austria. Understanding whether *D. apicalis* vector populations have adapted to specific conditions or may be able to spread over large areas will have implications for the control of psyllids as pests and potential emergence and spread of resistant genotypes.

## Acknowledgements

We thank the JIC Molecular Genetics, Research Computing, and Entomology platforms for their support, and Darren Heavens (Earlham Institute), Saleha Bakht (JIC), Elizabeth Hollwey (JIC, currently at ISTA, Austria) for useful discussions of experimental methods. We also thank Adrian Fox for management of the CaLiber consortium project. This work was funded by the CaLiber consortium (BB/T010851/1) through a grant from UK Research and Innovation (UKRI) under the Strategic Priorities Fund, in collaboration with the Biotechnology and Biological Sciences Research Council (BBSRC), and supported by the Department for Environment, Food and Rural Affairs (Defra) and the Scottish Government. Additional support was provided by the BBSRC Institute Strategy Programmes (BBS/E/J/000PR9797, BBS/E/J/000PR9798 and BBS/E/JI/230001B) awarded to the John Innes Centre (JIC), which is grant-aided by the John Innes Foundation.

## Contributions (please add any descriptions of contributions, and add other people as appropriate)

Thomas Heaven, Thomas Mathers, Sam T. Mugford and Saskia A. Hogenhout, JIC, design and implementation of the research, analysis of the results and writing of the manuscript. Jay K. Goldberg contributed to data analysis and writing the manuscript.

Fiona Highet, SASA, coordinated collection and shipment of psyllids from collaborators across Europe.

Jason Sumner-Kalkun and Jennifer Newton, SASA – collected samples from Scotland and reared colonies of psyllids for shipment to JIC

Saskia A. Hogenhout and Fiona Highet, Project and staff management.

Anne Nissinen-LUKE, Finland-reared colonies of psyllid and shipment to JIC, provided advise on conditions for psyllid rearing.

Anna Jordan and Victor Soria-Carrasco, developed methods and reared psyllid colonies. Christa Lethmayer, collected samples from Austria.

Lars-Arne Høgetveit, collected samples from Norway.

All authors contributed to and approved the final manuscript.

## Data Availability Statement

Raw reads used to generate our assemblies can be found on NCBI under BioProject accession number PRJNA1163903. BioSample and SRA accessions are detailed in Supplementary Table S6 (Supplementary Material online). Processed genomic datasets (assemblies, annotations, OrthoFinder/CAFE5 outputs, etc.) can be found on Zenodo at https://zenodo.org/records/13846379

## Supplementary Materials

**Table S1.** *Dyspersa apicalis* and *Dyspersa pallida* resequencing samples. Sampling location, method and date are given for each sample as well as sex where known.

**Table S2.** RNA-seq mapping statistics obtained from STAR alignments to our BRAKER annotations

**Table S3.** Genomes/Proteomes used for phylogeny construction, orthology analysis, and Computational Analysis of gene Family Evolution.

**Table S4.** Gene predictions using different methods and parameters from the genomes of *Dyspersa apicalis, Dyspersa pallida*, and *Trioza urticae*. Number of proteins and Omark and BUSCO results are given for each approach. The final annotations selected are highlighed in orange.

**Table S5.** Detailed KEGG enrichment results obtained from ClusterProfiler. These results are visualized in Figures S7-9.

**Table S6.** Data availability - NCBI - SRA Archive accession numbers.

**Figure S1.** Misassembled D. apicalis / *Ca*. C. ruddii sequence. A region of *D. apicalis* assembly with blast hits to the genome of *Ca*. C. ruddii was extracted and aligned to the *Ca*. C. ruddii reference genome (CP019943.1). An approximately 170 kb region had collinearity to the *Ca*. C. ruddii genome (A) has increased coverage (B), and as seen in the IGV view of “sorted.bam” track is flanked by zero and one × coverage positions (C).

**Figure S2.** Transposable element content of *Diaphorina citri* assemblies. GCA_024506325.2 (A), GCA_024506315.2 (B),and GCA_030643865.1 (C). Pie charts show TE content, different colours representing different TE superfamilies. Paired Kimura distance plots indicate the relative activity of different TE superfamilies in the genome.

**Figure S3.** Transposable element content of additional *Diaphorina citri* assemblies GCA_024506275.2 (A) and GCA_000475195.1 (B). Pie charts show TE content, different colours representing different TE superfamilies. Paired Kimura distance plots indicate the relative activity of different TE superfamilies in the genome.

**Figure S4.** Transposable element content of *Pachypsylla venusta* assemblies GCA_-12654025.1 (A) and GCA_000695645.2 (B). Pie charts show TE content, different colours representing different TE superfamilies. Paired Kimura distance plots indicate the relative activity of different TE superfamilies in the genome.

**Figure S5.** Transposable element content of *Bactericera cockerelli* assembly GCA_024516035.1. Pie charts show TE content, different colours representing different TE superfamilies. Paired Kimura distance plots indicate the relative activity of different TE superfamilies in the genome.

**Figure S6.** Phylogeny of *Candidatus* Carsonella ruddii samples from different psyllid hosts. Downloaded genomes are identified by their host species and GenBank accession ID, *Ca*. C. ruddii sequences generated as part of this study are also included.

**Figure S7.** Enriched KEGG terms in significantly expanded *Dyspersa apicalis* orthogroups.

**Figure S8.** Enriched KEGG terms in significantly expanded *Dyspersa pallida* orthogroups.

**Figure S9.** Enriched KEGG terms in significantly expanded *Dyspersa* orthogroups.

