## Supplemental Figures for "Chromosome-level Assemblies of Three *Candidatus* Liberibacter solanacearum Vectors: *Dyspersa apicalis* (Förster, 1848)*, Dyspersa pallida* (Burckhardt, 1986), and *Trioza urticae* (Linnaeus, 1758) (Hemiptera: Psylloidea)"

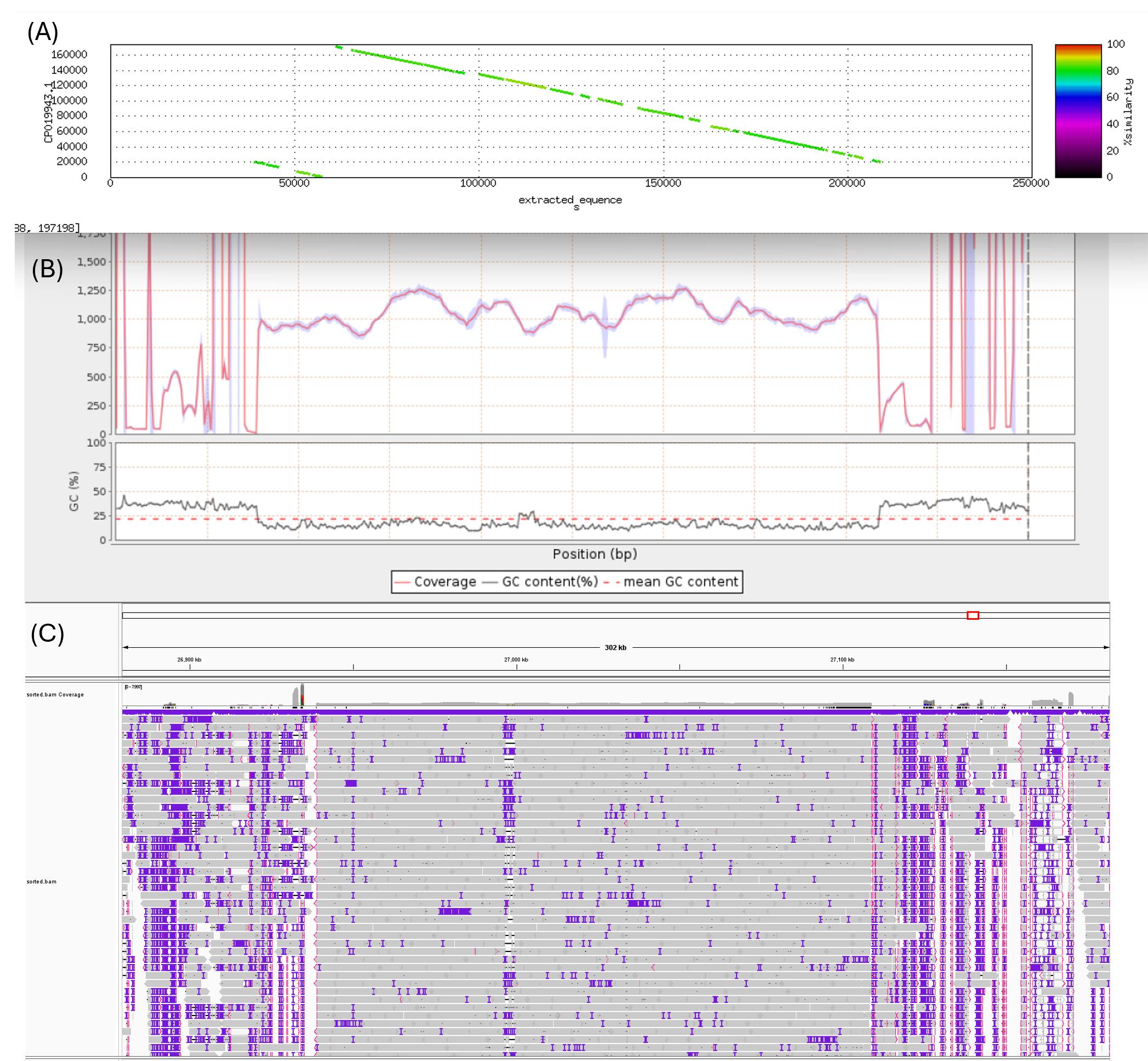

**Figure S1 – Misassembled *D. apicalis* / *Ca.* C. ruddii sequence**. A region of *D. apicalis* assembly with blast hits to the genome of *Ca.* C. ruddii was extracted and aligned to the *Ca.* C. ruddii reference genome (CP019943.1). An approximately 170 kb region had collinearity to the *Ca.* C. ruddii genome (A) has increased coverage (B), and as seen in the IGV view of “sorted.bam” track is flanked by zero and one × coverage positions (C).

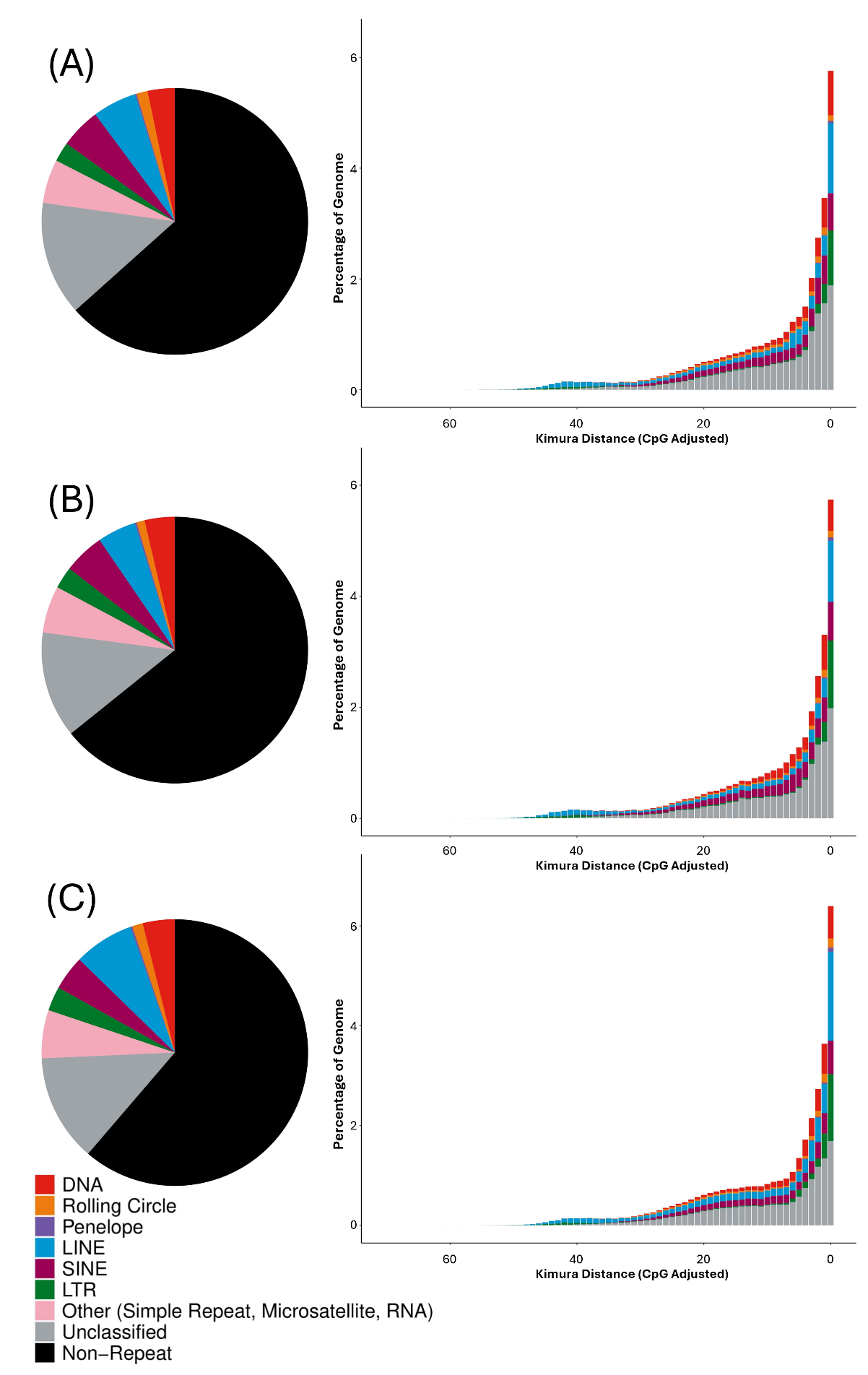
 **Figure S2 - Transposable element content of *Diaphorina citri*** assemblies GCA_024506325.2 **(A)**, GCA_024506315.2 **(B)**,and GCA_030643865.1 **(C)**. Pie charts show TE content, different colours representing different TE superfamilies. Paired Kimura distance plots indicate the relative activity of different TE superfamilies in the genome.

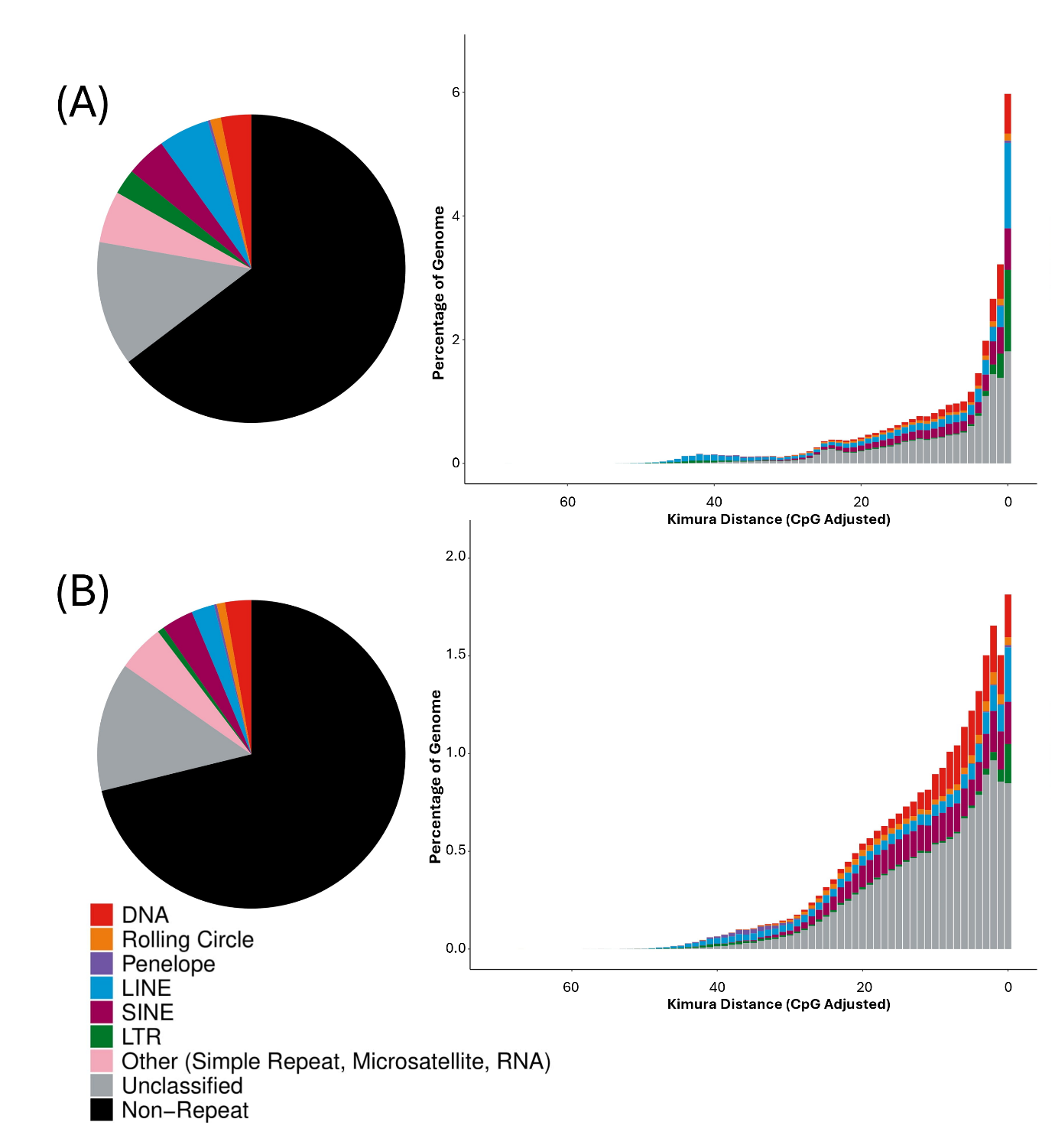

**Figure S3 - Transposable element content of additional *Diaphorina citri*** assemblies GCA_024506275.2 **(A)** and GCA_000475195.1 **(B)**. Pie charts show TE content, different colours representing different TE superfamilies. Paired Kimura distance plots indicate the relative activity of different TE superfamilies in the genome.

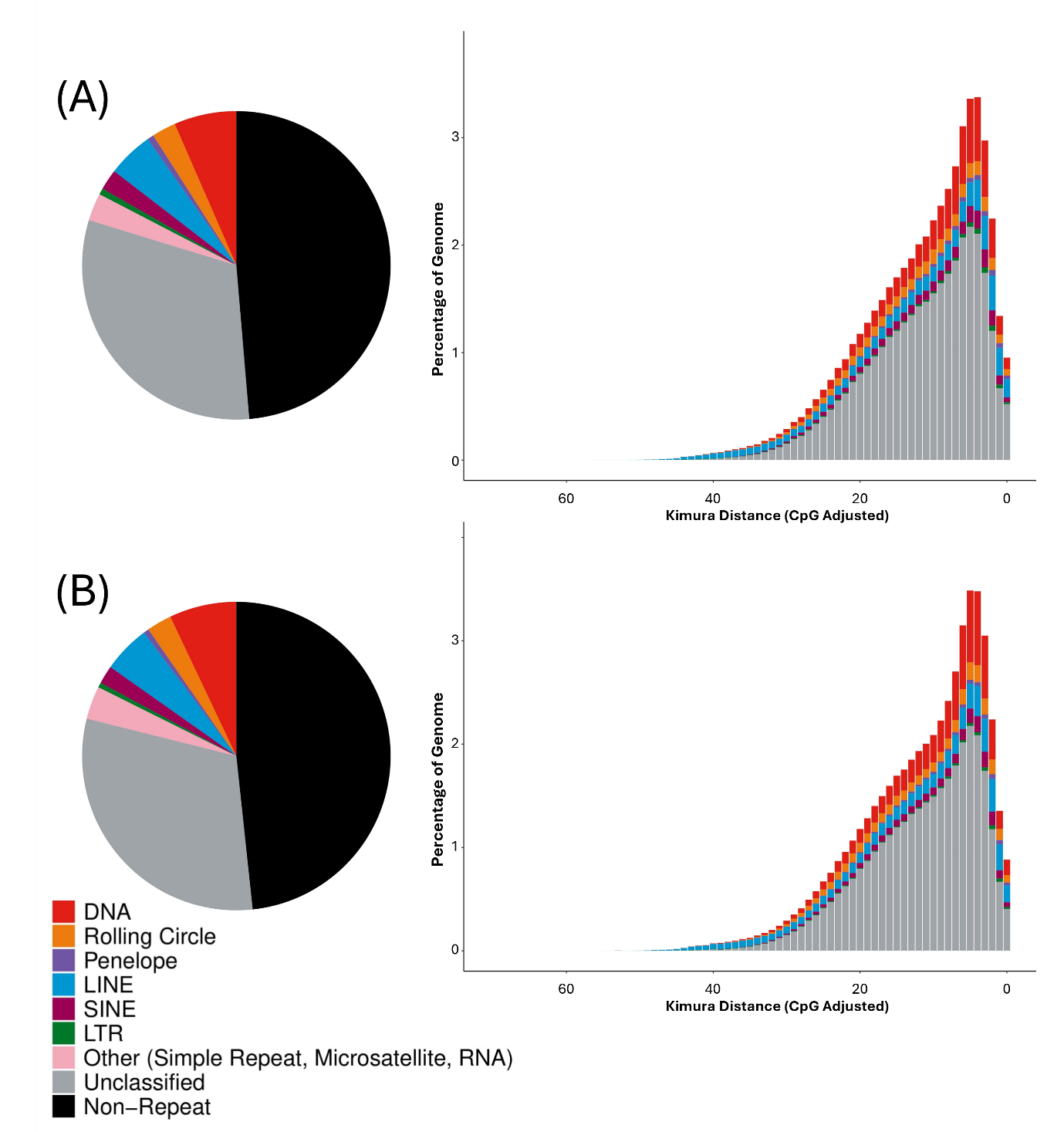

**Figure S4 - Transposable element content of *Pachypsylla venusta*** assemblies GCA_-12654025.1 **(A)** and GCA_000695645.2 **(B)**. Pie charts show TE content, different colours representing different TE superfamilies. Paired Kimura distance plots indicate the relative activity of different TE superfamilies in the genome.

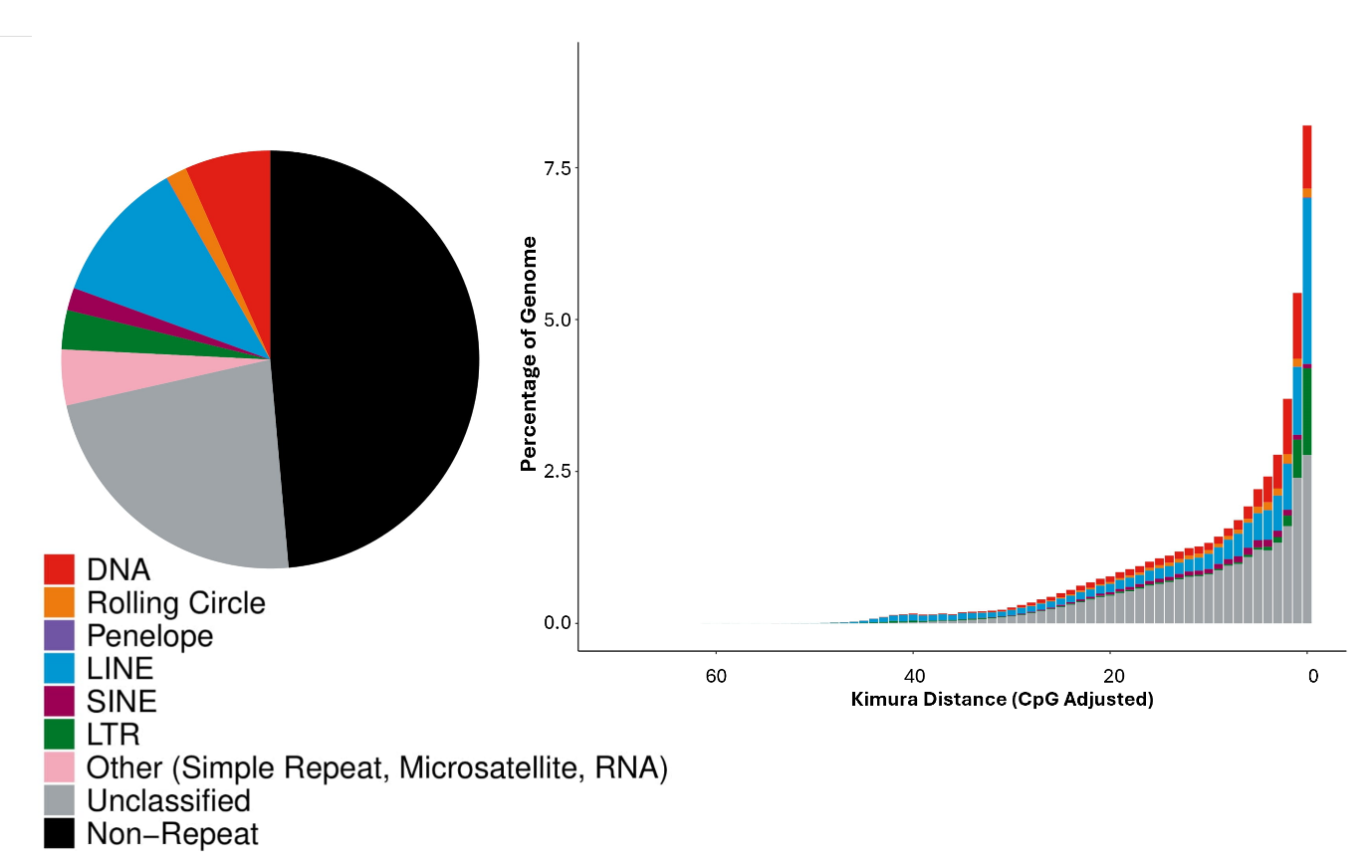

**Figure S5 - Transposable element content of *Bactericera cockerelli*** assembly GCA_024516035.1. Pie charts show TE content, different colours representing different TE superfamilies. Paired Kimura distance plots indicate the relative activity of different TE superfamilies in the genome.

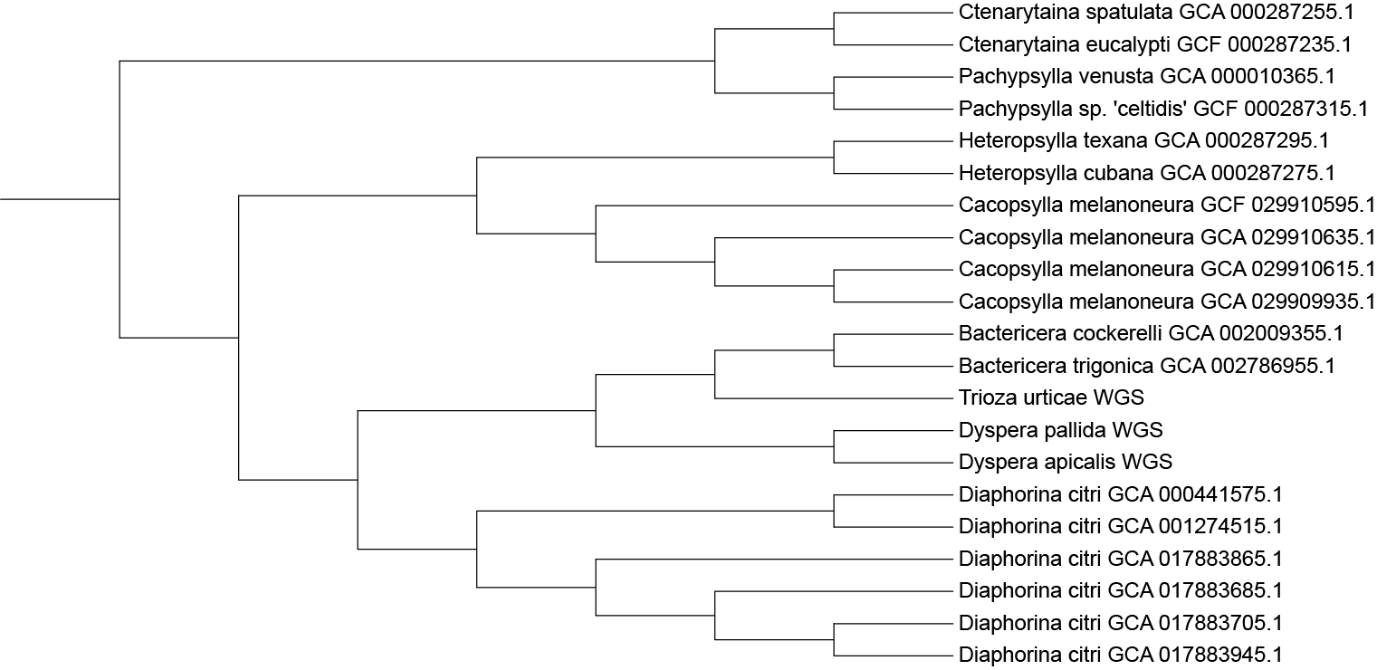

**Figure S6 – Phylogeny of *Candidatus* Carsonella ruddii samples from different psyllid hosts.** Downloaded genomes are identified by their host species and GenBank accession ID, *Ca.* C. ruddii sequences generated as part of this study are also included.

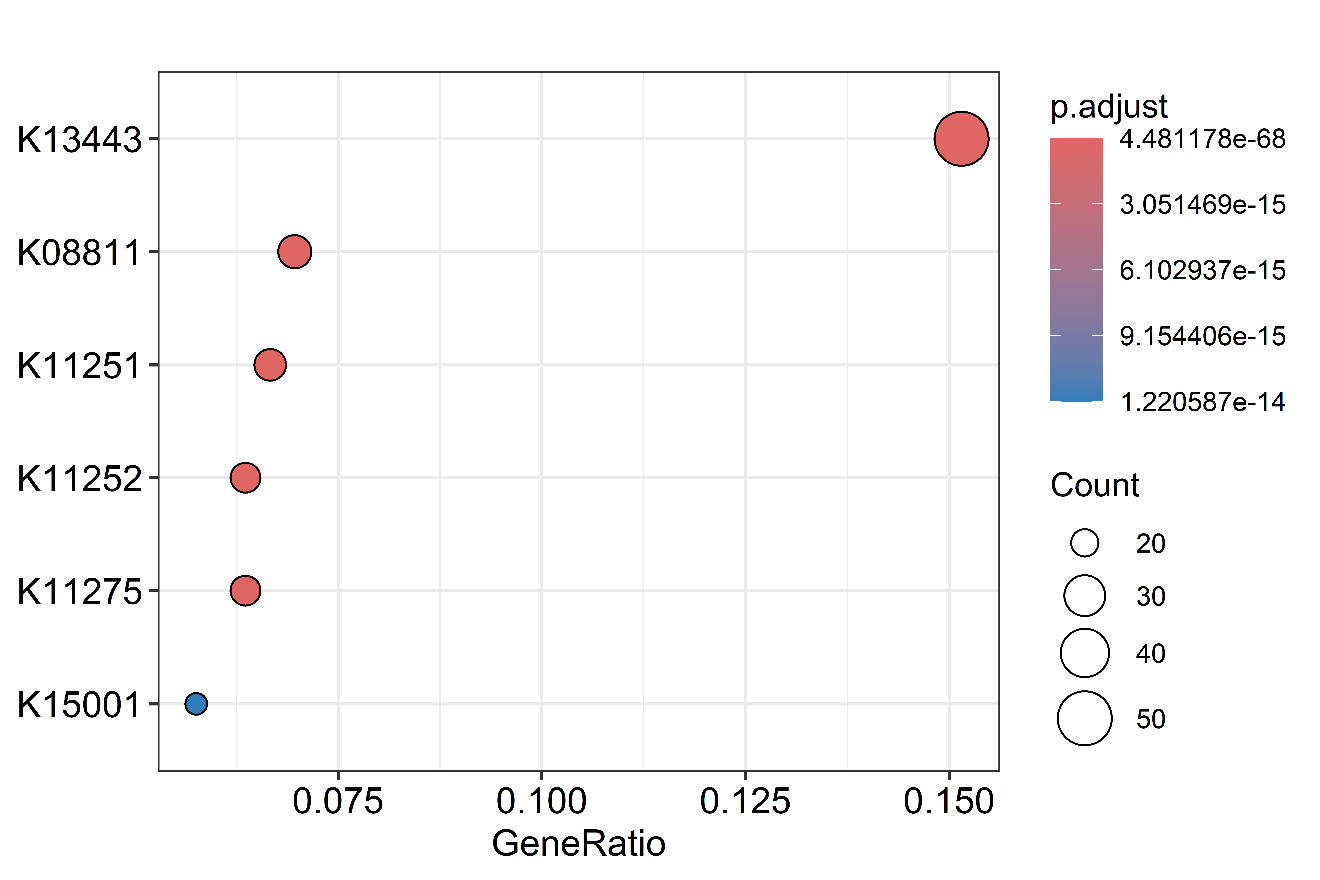

Figure S7 –**Enriched KEGG terms in significantly expanded *Dyspersa apicalis* orthogroups.**

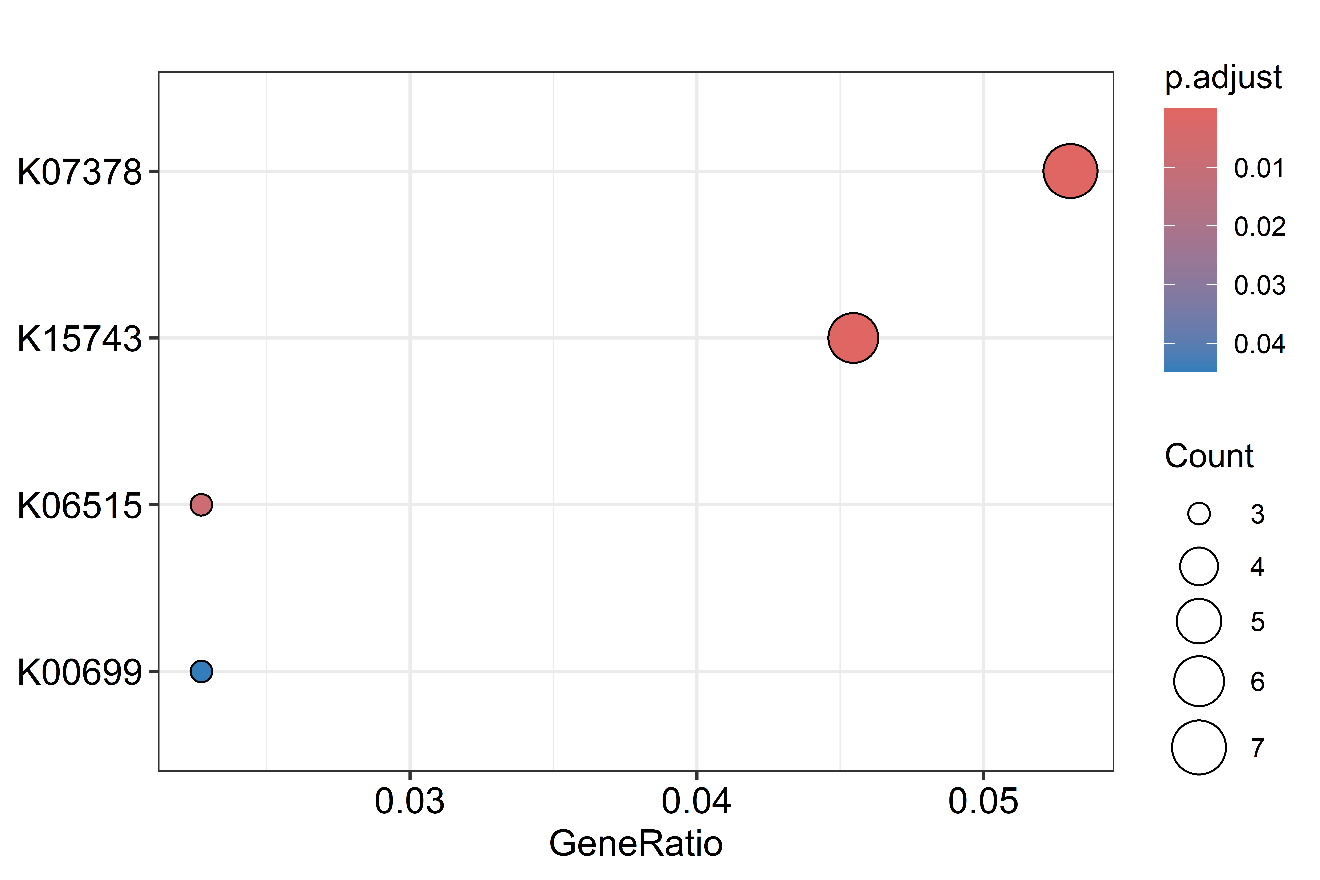

Figure S8 –**Enriched KEGG terms in significantly expanded *Dyspersa pallida* orthogroups.**

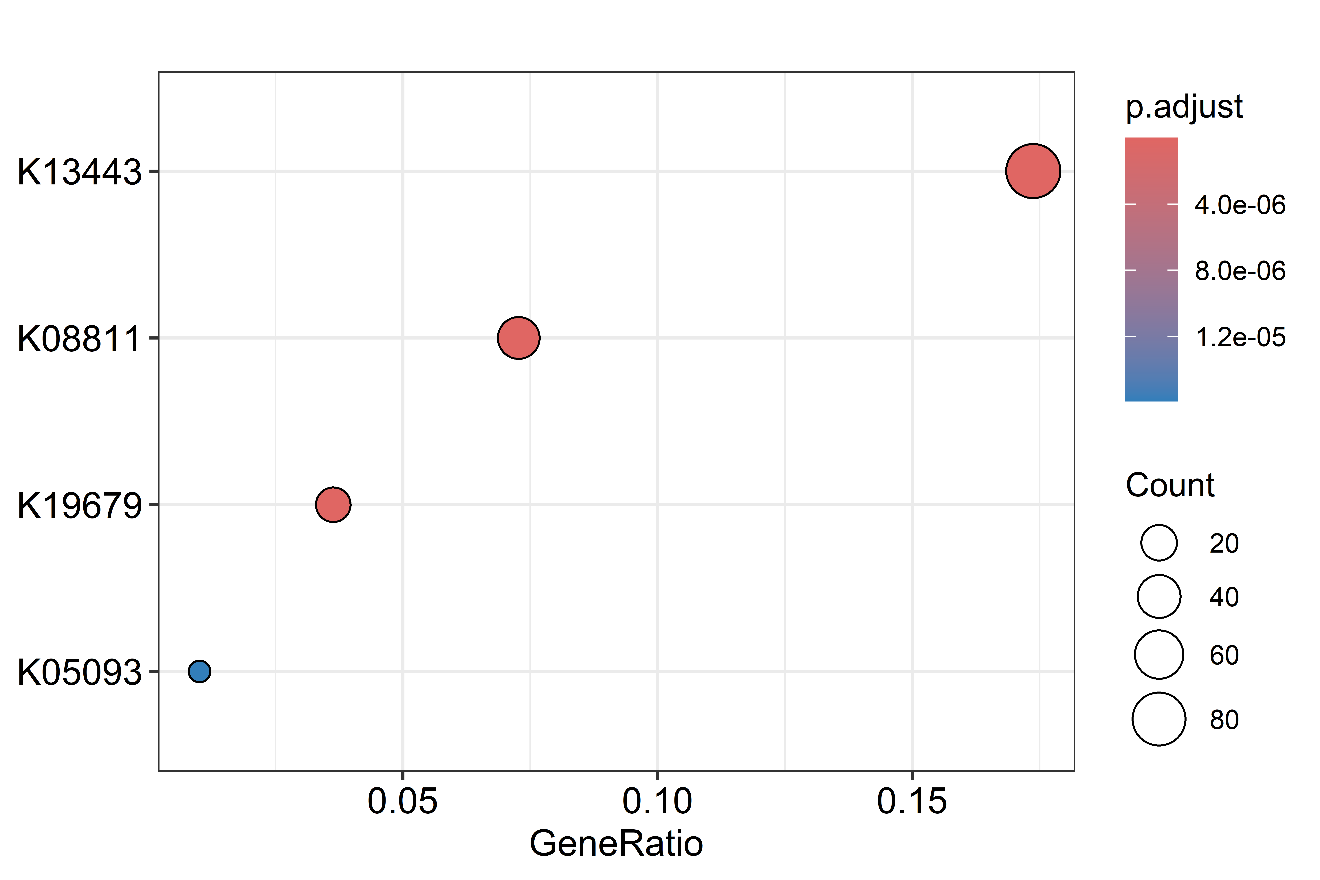

Figure S9 –**Enriched KEGG terms in significantly expanded *Dyspersa* orthogroups.**
